# Structure of the human SAGA coactivator complex: The divergent architecture of human SAGA allows modular coordination of transcription activation and co-transcriptional splicing

**DOI:** 10.1101/2021.02.08.430339

**Authors:** Dominik A. Herbst, Meagan N. Esbin, Robert K. Louder, Claire Dugast-Darzacq, Gina M. Dailey, Qianglin Fang, Xavier Darzacq, Robert Tjian, Eva Nogales

## Abstract

Human SAGA is an essential co-activator complex that regulates gene expression by interacting with enhancer-bound activators, recruiting transcriptional machinery, and modifying chromatin near promoters. Subunit variations and the metazoan-specific requirement of SAGA in development hinted at unique structural features of the human complex. Our 2.9 Å structure of human SAGA reveals intertwined functional modules flexibly connected to a core that distinctively integrates mammalian paralogs, incorporates U2 splicing subunits, and features a unique interface between the core and the activator-binding TRRAP. Our structure sheds light on unique roles and regulation of human coactivators with implications for transcription and splicing that have relevance in genetic diseases and cancer.

---

In eukaryotes, expression of protein-coding genes involves the integration of cellular signals, chromatin modification, and assembly of the transcription pre-initiation complex (PIC) that loads RNA polymerase II onto the core promoter (*1, 2*). PIC formation is initiated by recruitment of the TATA-binding protein (TBP) to the promoter by protein complexes containing TBP-associated factors (TAFs) that interact with activators, promoter DNA, and/or chromatin (*3*). Throughout evolution, these TAFs partitioned and specialized into two distinct coactivator complexes in eukaryotes, the general transcription factor TFIID and the Spt-Ada-Gcn5 acetyltransferase (SAGA) complex (*3, 4*). Structural studies on human TFIID (hTFIID) have elucidated its molecular function as a chaperone for TBP deposition and initiation of PIC assembly (*5*). Studies of TFIID and SAGA in yeast highlight these TAF-containing complexes as global coactivators of transcription, but their unique or overlapping roles in human gene regulation are still not well understood (*6-8*). SAGA has been functionally implicated in a multitude of cellular pathways, from serving as a regulatory hub in transcription, to its involvement in cell proliferation (*9*). With more than a megadalton in size, SAGA contains several distinct functional modules (Fig. 1A): a core module that forms a structural scaffold of histone-fold (HF)-containing TAFs; a TRRAP (Transformation/Transcription domain Associated Protein) module that contains a pseudo-PI3 kinase (ΨPI3K) (*10*) and binds activators such as c-Myc, E2F, and p53 (*10, 11*); a histone acetyltransferase (HAT) module that deposits H3K9ac and H3K14ac at promoters of active genes (*7, 12*); and a deubiquitinase (DUB) module that removes H2B K120 ubiquitination from active gene bodies (*7, 13*).

**Fig. 1.**
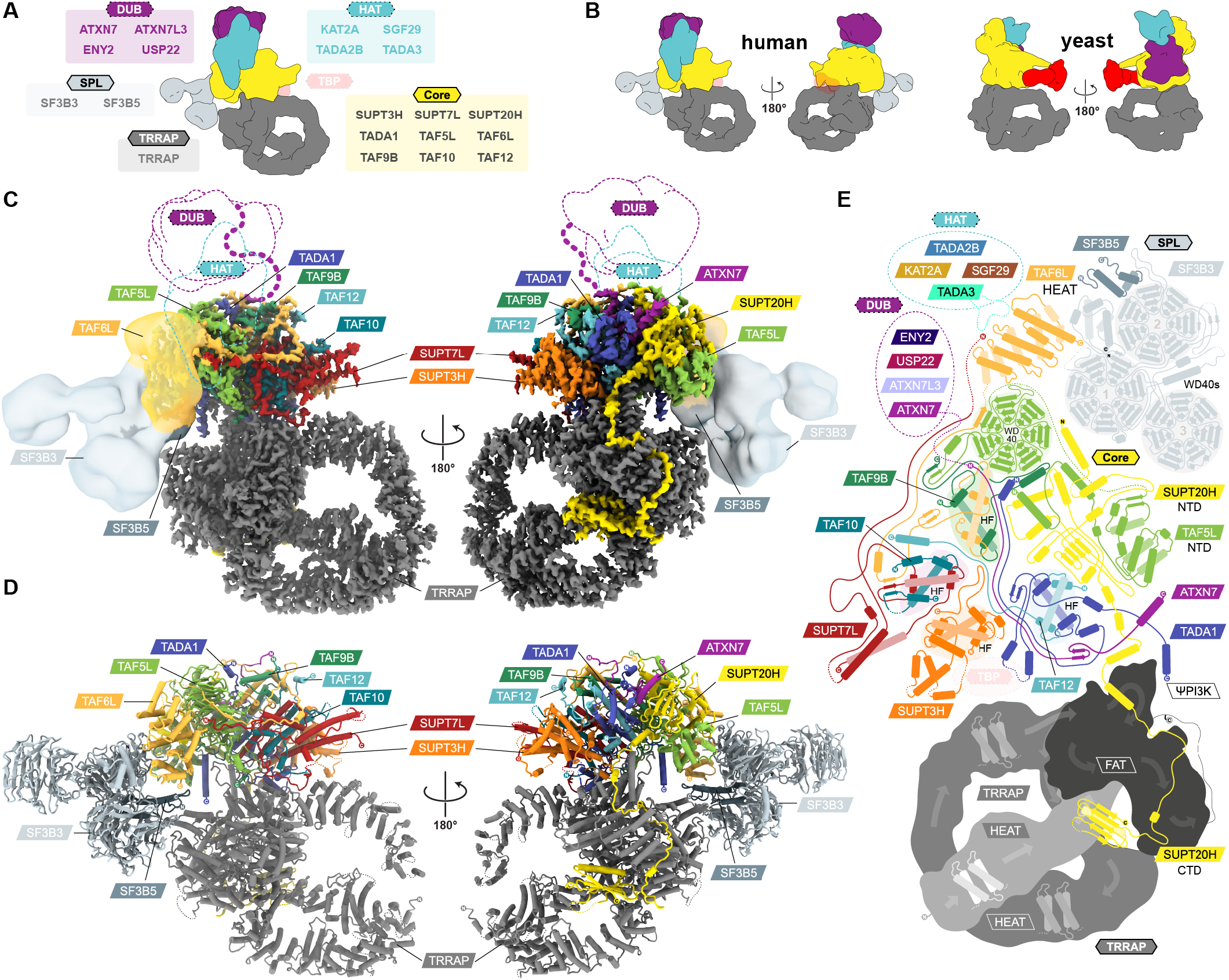
Architecture of the human SAGA complex. (**A**) Schematic of the modular architecture and subunit composition of hSAGA. The putative position of TBP is indicated with a transparent surface, based on ySAGA. (**B**) Comparison of the modular organization of hSAGA (left) and ySAGA (*16*) (right). Modules are colored according to panel A, with yeast modules colored according to their human homolog. (**C**) Hybrid map of hSAGA. The high-resolution cryo-EM map shows the best resolved core and TRRAP modules as a solid surface (contoured at 7.0σ), and the negative stain map for the TAF6L HEAT domain and the SPL module shown as a transparent surface (contoured at 6.1σ). The expected locations of DUB and HAT are indicated as dotted outlines, and the DUB module connection to the core via ATXN7 is shown as a purple dashed line. Subunits are colored as indicated by the labels. (**D**) Atomic model of hSAGA. (**E**) Topology map of hSAGA subunits grouped by modules (subunits are not drawn to scale; see also Fig. S5). Color schemes for modules and subunits are consistent throughout all figures.

Initial structural studies on SAGA were limited to isolated regions or domain fragments (*14*). More recent studies on the complete yeast SAGA (ySAGA) complex have revealed its modular organization, and structural details for two of its modules (*15-17*).

While vertebrate SAGA is highly conserved (∼95-58% sequence identity (seqID)), the conservation with yeast drops dramatically (∼18% seqID) (Table S1) and numerous domain insertions and deletions as well as gene duplication, have led to subfunctionalization of human SAGA (hSAGA) subunits (*4*). Substantial evolutionary differences are further indicated by the lack of a metazoan homolog to one TBP-binding subunit in ySAGA (Spt8), and by the incorporation of the splicing factor subunits SF3B3 and SF3B5 within the metazoan complex (*18*), which gives rise to a fifth, splicing module (SPL) in metazoan SAGA. Lastly, while SAGA is not essential for viability in yeast, it is strictly required for metazoan development, and disruption of hSAGA can lead to broad pathogenic effects, including neurodegenerative disease and cancer (*9*). The difference in its essentiality, as well as the presence of specialized subunit paralogues and gained interaction partners indicate that some aspects of assembly and function will be clearly distinct between the human and yeast SAGA complexes. In this study, we sought to elucidate the architecture of hSAGA to examine the possible functional implications of its conserved and distinct features.

## Architecture of hSAGA

Human SAGA is a 1.4 MDa protein complex composed of 20 subunits. Low endogenous expression, dynamic composition, and fragile complex assembly have challenged the structural characterization of full hSAGA. In order to obtain intact endogenous hSAGA for structural studies, we genome edited a HeLa cell line using CRISPR-Cas9 to express Halo-FLAG-tagged chromosomal *SUPT7L*. Affinity purification yielded a few picomoles of pure complex from 30 L of HeLa cells. We validated the presence of all subunits within our hSAGA sample using Western blotting and mass spectrometry (MS) (Fig. S1, Table S2, S3, supplementary methods). While no significant amount of endogenous TBP was co-purified with the complex (Fig. S1), a significant amount of the SPL module subunits SF3B3 and SF3B5 were found complexed with hSAGA (Table S2, S3).

We first examined hSAGA by negative stain electron microscopy. Early 3D classification revealed a central core sitting atop the distinct cradle-shaped TRRAP module (Fig. S2A). We also observed a highly variable Y-shaped density proximal to the TRRAP cradle, indicating a flexible tethering to the core. Final classification resulted in reconstructions with and without this flexible region, with the one including it refining to 19 Å resolution (Fig. S2B-C). A comparison of hSAGA and ySAGA shows clear architectural differences concerning the relative positions of the core and TRRAP modules (see below), and the additional Y-shaped density in hSAGA that we were able to unambiguously assign to the metazoan-specific splicing module (Fig. 1B, Table S4).

Using cryo-electron microscopy (cryo-EM) we obtained a reconstruction at an overall resolution of 2.9 Å for the best ordered regions of hSAGA, the TRRAP and core modules, (Fig. 1C, Fig. S3A-E, Table S4) that allowed us to build atomic models of these regions (Fig. 1D, Fig. S3F-G, Table S4). A homology model of the TAF6L HEAT (Huntingtin, elongation factor 3 (EF3), protein phosphatase 2A (PP2A), and the yeast kinase TOR1) domain (*5*) and the crystal structure of SF3B3/SF3B5 (*19*) within the splicing module were docked into their corresponding positions in the density map, which were observed only in the lower resolution, negative stain reconstruction (Fig. S2D). In the higher resolution map, fuzzy density for the TAF6L HEAT domain can be observed after LocSpiral filtering (Fig. S4A), demonstrating the flexible character of this region. The superposition of the two density maps revealed the architectural integration of the SPL module within hSAGA (Fig. 1C-E, Fig. S5).

Although all HAT and DUB subunits were present in our sample, they were not resolved in our structural analysis, either due to flexible tethering or a more dynamic and labile attachment. Nevertheless, we expect the DUB and HAT modules in hSAGA to be located at similar positions with respect to the core as seen in ySAGA. Concerning the HAT, both the yeast and human complexes exhibit the same relative location for the N-terminal region of SUPT7L within the core module, which binds to the Ada3/TADA3 subunit of the HAT module in yeast (*16*) (Fig. S6A-C). For the DUB, we could model a segment of the DUB subunit ATXN7 that connects this module with the rest of the complex in a similar location as in yeast (Fig. S6D-F). Notably, both the HAT and DUB modules were only resolved at low resolution in the ySAGA structures, highlighting their intrinsic flexibility across species (*15, 16*). Structural plasticity of the HAT and DUB modules may be critical for SAGA’s function since the HAT module mainly acetylates promoter chromatin while the DUB acts within gene bodies. Some flexibility may allow the modules to perform these cellular functions independently, spatially and/or temporally (*7, 20*).

### The core module

The structure of hSAGA is organized around a core module that consists of nine subunits (Fig. 1) and connects to the rest of the functional modules within the complex. Seven subunits of the core contain histone fold (HF) domains (TAF6L, TAF9B, TAF10, TAF12, SUPT7L, TADA1, and SUPT3H, which contains two HFs) (Fig. 1E, 2A-B, Fig. S5) and assemble into a distorted pseudo-octamer (Fig. 2C), as also observed in ySAGA (*15, 16*) and in both human and yeast TFIID (*5, 21*). The two HF domains of SUPT3H form an H2A/H2B-like dimer, and in ySAGA they have been shown to bind TBP (*16*) (Fig. 2B). In hTFIID, TBP is bound by TAF11/13 in the same relative position (*5, 22*). On the opposite side of the octamer core, TAF6L binds centrally via the C-terminal end of helix ∝2 of its HF domain to the seven bladed WD40 propeller of TAF5L (Fig. 2A). This domain of TAF5L distorts the HF pseudo-octamer and organizes the core by acting as a platform for tethering neighboring domains that determine the location of the SPL and HAT modules. In the ySAGA structure, the Ada3 (human TADA3) subunit of the HAT module binds to the convex surface of the C-terminal TAF6L HEAT repeat domain, which we observe for the hSAGA complex in the negative stain analysis, and that likely also acts as the tether for the hSAGA HAT module (Fig. S6A-C, 1C, 1E). The increased flexibility in this region in the human complex is likely due to a lack of stabilization by the TAF5L N-terminal domain (NTD), which is rotated (relative to the TAF5L WD40 propeller) away from the TAF6L HEAT domain and latched in place by SUPT20H (Fig. 3A). This leaves the concave HEAT domain interface accessible for binding of the SPL module. In ySAGA, the corresponding Taf5 NTD is rotated -59° and binds to the same concave surface of the Taf6 HEAT domain (Fig. 3B). The yeast Taf5-Taf6 interaction is further stabilized by the Taf5 C-terminal NTD linker, which wraps around the side of the Taf6 HEAT domain (*15, 16*). Different orientations of the TAF5 NTD have also been observed in TFIID and are a crucial marker for a divergent architecture of that complex (*5, 21*) (Fig. 3C).

**Fig. 2.**
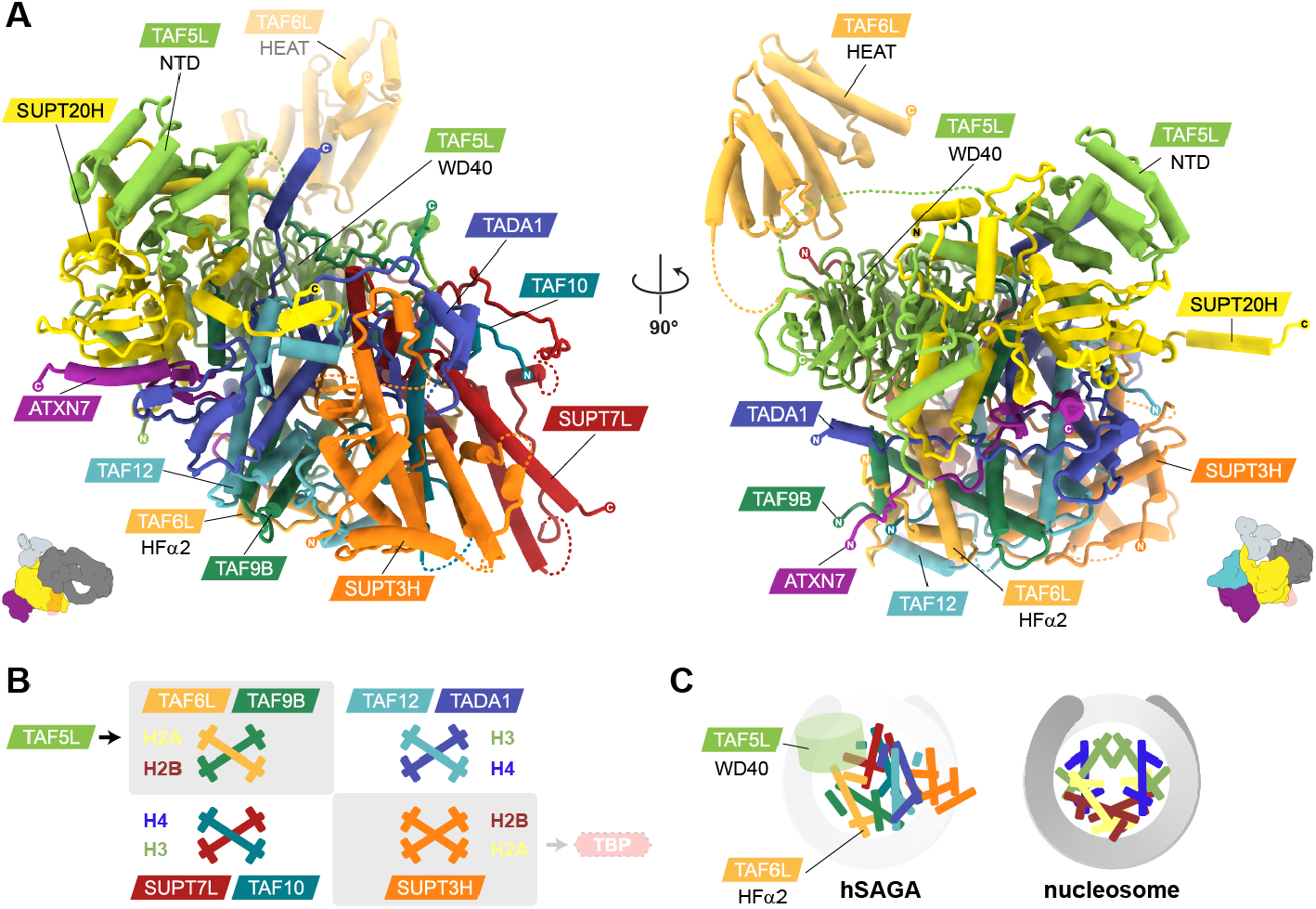
Organization of the distorted histone fold core module. (**A**) Orthogonal views of the atomic model of the pseudo-octamer formed by the HF-containing subunits in the hSAGA core, with the laterally attached TAF6L HEAT and TAF5L NTD domains. Relative orientations with respect those in Fig. 1A/B are indicated by the inset. The C-terminal end of the ∝2 helix of the TAF6L HF binds to the center of the WD40 propeller of TAF5L. (**B**) Comparison of the HF core organization in hSAGA with the nucleosome. HF dimers are shown in boxes, colored based on hSAGA subunits, with the relative subunit locations occupied by corresponding histones in the nucleosome indicated on the side, as well as the contact with the TAF5L WD40 propeller and the potential interaction with TBP. SUPT3H contains an intramolecular dimer of HFs. (**C**) Schematic of the relative locations of the distorted HF octamer helices in the hSAGA core and in the nucleosome. In hSAGA the distortion creates a gap that is occupied by the TAF5L WD40 propeller.

**Fig. 3.**
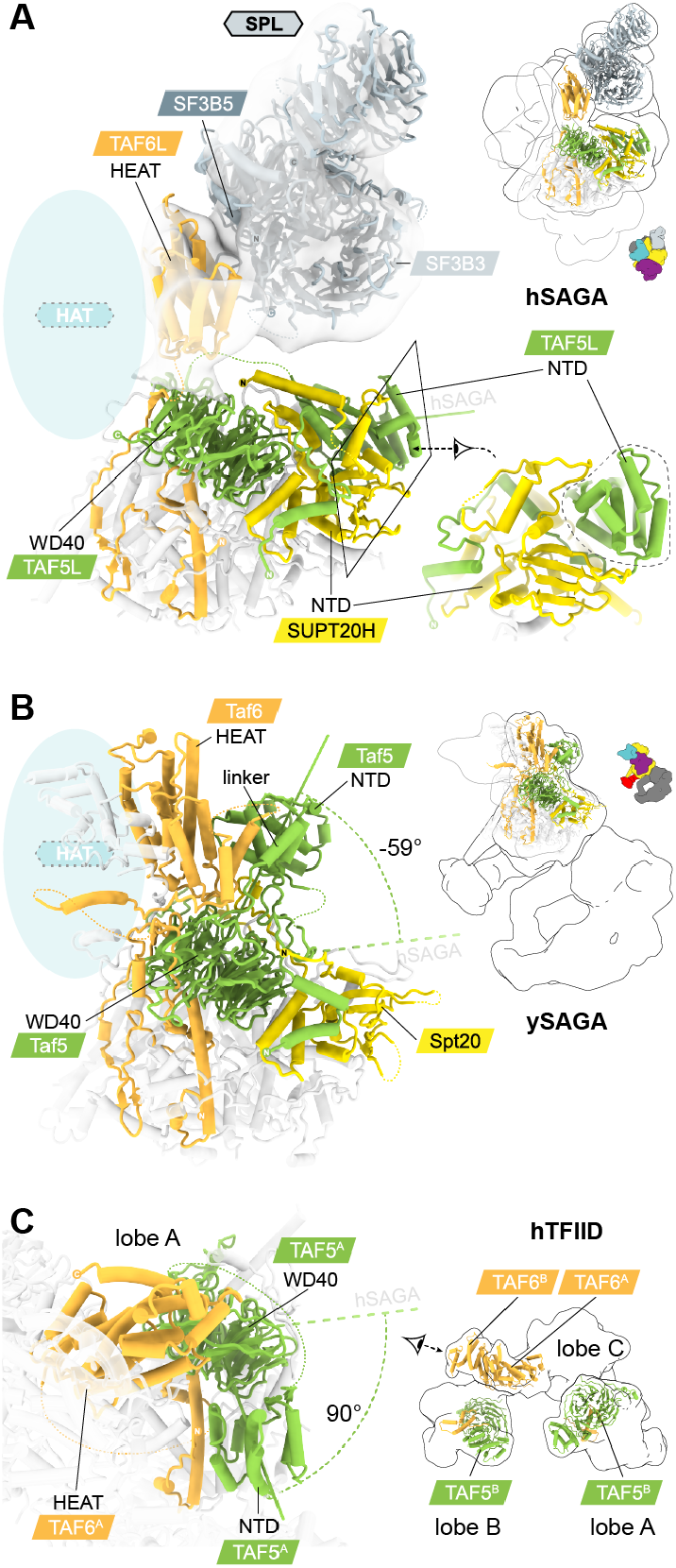
The HAT and SPL modules are integrated by TAF6L, TAF5L, and SUPT20H. (**A, B**) The (concave) surface of the TAF6L HEAT domain that tethers the SPL module in hSAGA (EM map for both shown in transparent and contoured at 6.1σ) (**A**), is engaged with the Taf5 NTD in ySAGA (PDB: 6T9K) (**B**). The opposite (convex) surface tethers the HAT module in both complexes (transparent blue oval). The relative location of the depicted region in each complex is indicated in the outlines on the right. Relative locations of all modules are indicated in the colored insets. In hSAGA, the SUPT20H NTD latches the TAF5L NTD in place (A) (a close-up view from a different angle is shown on the right). (**C**) Comparison with lobe A of hTFIID. TFIID contains two copies of TAF5 and TAF6, with one TAF5 located in lobe A and the other one in lobe B, and with the two TAF6 HEAT domains in lobe C (shown on the right). Compared to SAGA, the HEAT domains and NTDs adopt different relative positions with respect to the WD40 domains, leading to divergent architectures. NTD angles relative to hSAGA are indicated in all panels.

The subunit with the largest interface with the rest of the complex is SUPT20H (approx. 12,000 Å^2^), which forms a clamp-like scaffold within hSAGA (Fig. 4A). Several studies point to the central role of SUPT20H in complex assembly and module association, particularly for hSAGA. Deletion of Spt20, the homolog of SUPT20H in *S. pombe*, compromised incorporation of Tra1 (homolog of human TRRAP) and decreased DUB module association, but had little effect on the SAGA core (*23*). Markedly, depletion of SUPT20H in HeLa cells resulted in a large decrease of observable core and HAT SAGA subunits in ATXN7L3 pulldowns (*24*).

**Fig. 4.**
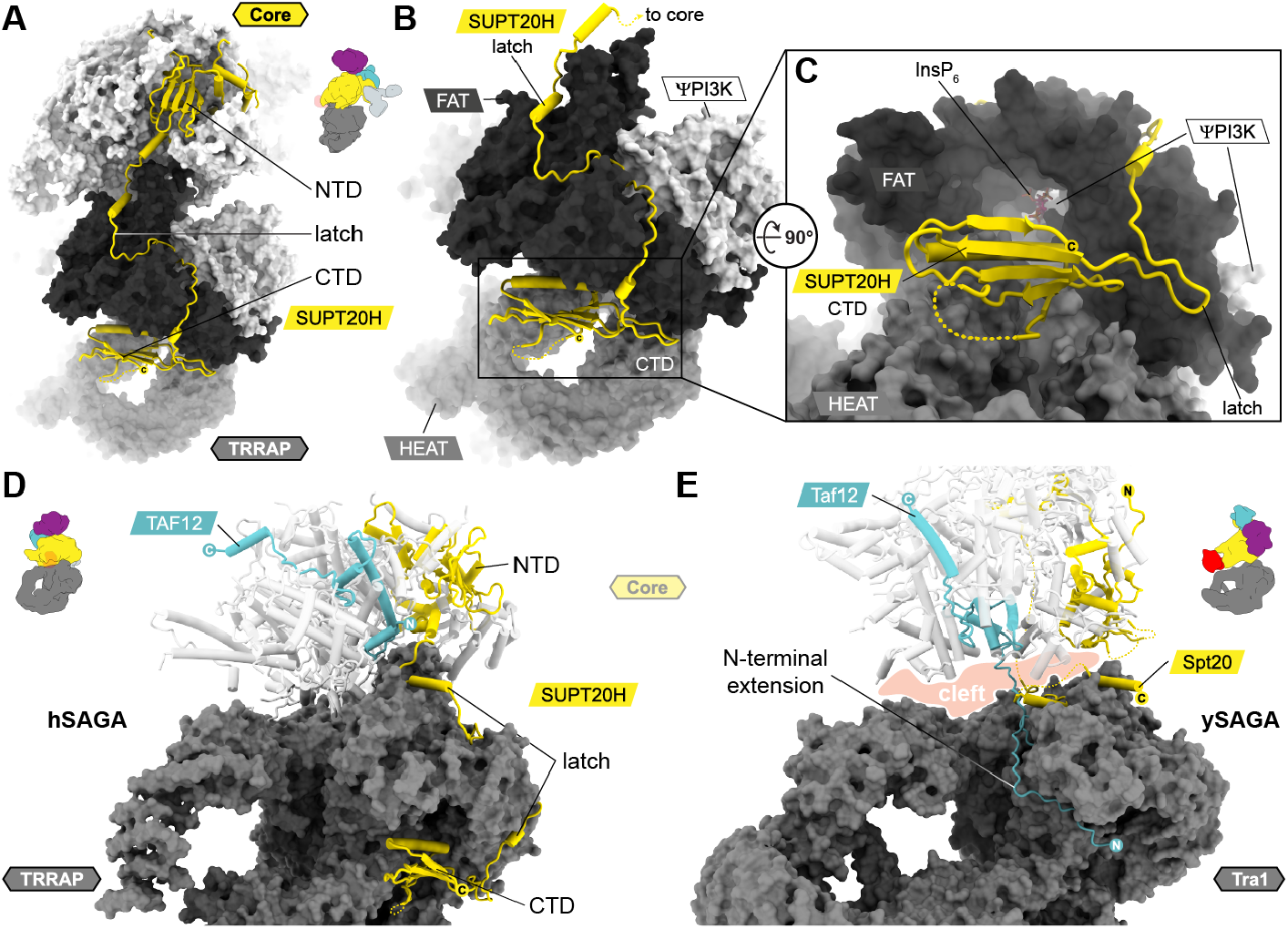
SUPT20H latches core and TRRAP together and contains a previously uncharacterized C-terminal domain. (**A**) SUPT20H contains two domains that bind to the core and TRRAP module, respectively, and are connected via a linker, the “latch”, that runs along the back of hSAGA. The relative orientation of the full complex is indicated by the colored inset. (**B**) Close-up view of the TRRAP surface, with TRRAP domains colored in different shades of grey. SUPT20H runs along a surface groove around the TRRAP FAT domain and terminates in a CTD. (**C**) The SUPT20H CTD is located at the entrance to a tunnel below the FAT domain that binds InsP_6_ (see Fig. S10). (**D, E**) In hSAGA the SUPT20H latch wraps around the outer surface of the TRRAP FAT domain (D). In yeast, Spt20 does not extent into the Tra1 module. Instead, Taf12 has an N-terminal extension that wraps around the inner surface of the Tra1 FAT domain (PDB 6T9I)(E). The core module is shown in cartoon representation. In contrast to hSAGA (D), ySAGA features a cleft between the Tra1 and core module (indicated in light red) (E).

In the core of hSAGA, the SUPT20H NTD holds the TAF5L NTD away from the TAF6L HEAT domain (Fig. 3A) allowing for docking of SF3B3 and tethering the N-terminus of the ATXN7 DUB anchor to the core (Fig. 2A), thus playing a critical role in connecting the DUB and (indirectly) the SPL modules to the core. Following the SUPT20H NTD is a long linker, hereafter referred to as the latch (Fig. 4A), that runs along the surface of the core, crosses the gap to the TRRAP, continues to wrap along the surface of the TRRAP FAT domain and terminates with an unpredicted C-terminal domain (CTD) in the cleft below the FAT and central TRRAP HEAT repeat (Fig. 4B). The CTD folds into a five stranded antiparallel beta sheet with an alpha helix parallel to the sheet that connects the two C-terminal outer strands (Fig. 4C, 1C-E). Neither the sequence of the SUPT20H latch, nor the CTD are conserved in ySAGA (Fig. S7A, B), indicating that the CTD is unique to the metazoan complex and explaining different functional relevance of this subunit in yeast and human assembly. Of notice, the N-terminus of ySAGA Taf12 emerges out of the region that is occupied by the SUPT20H CTD in hSAGA and wraps around the opposite side of the Tra1 FAT domain (Fig. 4D, E, S7C, D). Human and metazoan TAF12 have a much shorter N-terminus, contain gaps in the sequence corresponding to the Tra1 interface in yeast, and they contact TRRAP at a different location (Fig. S7C, D).

### Human SAGA and TBP

In ySAGA, TBP is bound via the core subunit Spt3 (homolog of the human SUPT3H) and by the yeast-specific Spt8 subunit, defining a separate TBP binding module that does not exist in humans (*16, 18*). Spt3 binds one end of the saddle-shaped TBP, while the other end is stabilized by binding to the beta propeller of Spt8. Spt8 itself contributes significantly to the interaction with TBP and has been shown to be sufficient to bind TBP alone, whereas Spt3 is not (*25*). Spt8 is absent in metazoans, and, to the best of our knowledge, biochemical evidence of TBP binding by the human homolog SUPT3H or the hSAGA complex is still lacking. Consistent with previous studies (*15, 16, 26-28*), our endogenous hSAGA sample did not co-purify with detectable levels of endogenous TBP (Fig. S1B). Interestingly, further analysis of our cryo-EM structure (see Materials and Methods) improved map quality of the region where SUPT3H, SUPT7L, TADA1, and TRRAP meet, which includes the region where TBP binds to in ySAGA (*16*), and suggested alternative main chain conformations that could not be sorted out by classification (Fig. 1E, Fig. S4B, C, Fig S5). To further examine TBP binding, we studied hSAGA by cryo-EM in the presence of added excess human TBP, but no additional density was observed (see Materials and Methods). These results, together with the heterogeneity in the cleft between SUPT3H, TADA1, and TRRAP, likely indicate a unique, highly dynamic mode of TBP binding by hSAGA, unlike that for the TFIID or the ySAGA complexes. The absence of Spt8 and the well characterized interaction of c-Myc with TBP (*29*) as well as TRRAP (*10, 30*) by separate c-Myc regions, might indicate that activators could play a role in TBP recruitment to metazoan SAGA. The ySAGA structure indicated that DNA binding by TBP is sterically hindered by Tra1 (homolog of human TRRAP) (*16*). Because of the distinct angle and connection of human TRRAP with the hSAGA core, the positioning of TBP within the hSAGA architecture may also have different consequences on TBP-DNA binding or on PIC assembly.

### TRRAP module

The characteristic TRRAP module has a tripartite HEAT repeat organization, consisting of a central N-terminal repeat and a circular cradle, followed by a FAT (Focal adhesion kinase targeting) domain and a ΨPI3K that connects to the core module (Fig. 1E, S5) and is embedded in the HEAT repeats, as also found in ySAGA, yeast NuA4 and its metazoan counterpart Tip60 (*15, 16, 31*), and in active kinases such as mTOR (*32, 33*), DNA-PKcs (*34*), and ATM (*35*) (Fig. S8A-E). However, the SAGA ΨPI3K lacks the canonical active site residues for catalysis (*10, 36*) (Fig. S8F). Our structure shows that the first residue of the TRRAP activation loop (Y3698), corresponding to the aspartate in the canonical PI3K DFG motif in active kinases (*10*), adopts an unusual cis-peptide bond that is well supported by the density map (Fig. S8G). Such non-proline geometry outliers often have a specific function within active sites (*37*), and its position in our structure, together with the high evolutionary conservation of the ΨPI3K, raises the question of whether the inactive kinase domain might have a different and so far undiscovered function, as observed for various pseudokinases in signaling and disease (*38*).

The most significant structural disparity between hSAGA and ySAGA is a fundamentally different interface between TRRAP and the core, which in hSAGA is rotated by 75° around the core subunit SUPT3H (Fig. 5A, S9A-C). In yeast, this interface is formed by Taf12, Spt20, Spt3, and partially by Ada1 (*15, 16*) and is separated from the core by a large cleft (Fig. 4E, S9D-I) that does not exist in hSAGA and presumably increases flexibility between the modules (Fig. S9C, F, I) (*17, 39*). In hSAGA, all core subunits except for TAF6L and ATXN7 are involved in establishing the interface to the TRRAP (Fig. 5B, C). While the central interface (Fig. 5B, S9B,C), corresponding to the main footprint of the core on TRRAP/Tra1, has a similar size (approx. 3,500 Å^2^), both complexes rely on additional stabilization by unique extensions of either Taf12 (Fig. 4E, S7C, S9F,I,J) or SUPT20H (Fig. 4D, S7A, S9C), which doubles the interface in hSAGA (approx. 7,070 Å^2^) and makes it 64% larger than that of ySAGA (approx. 4,300 Å^2^) (Fig. S9J). In yeast, Tra1 is shared between ySAGA and NuA4 (*40, Patel, Zukin and Nogales, in preparation*) and connects to the rest of the complex via a similar interface region on Tra1, suggesting that the TRRAP interface in hSAGA might also be relevant for the related metazoan Tip60 complex. Despite the extensive interface between the core and TRRAP modules in hSAGA, their connection is not rigid, as indicated by the relative motions observed between them upon analysis of a multi-body refinement (Fig. S4D) (see Material and Methods).

**Fig. 5.**
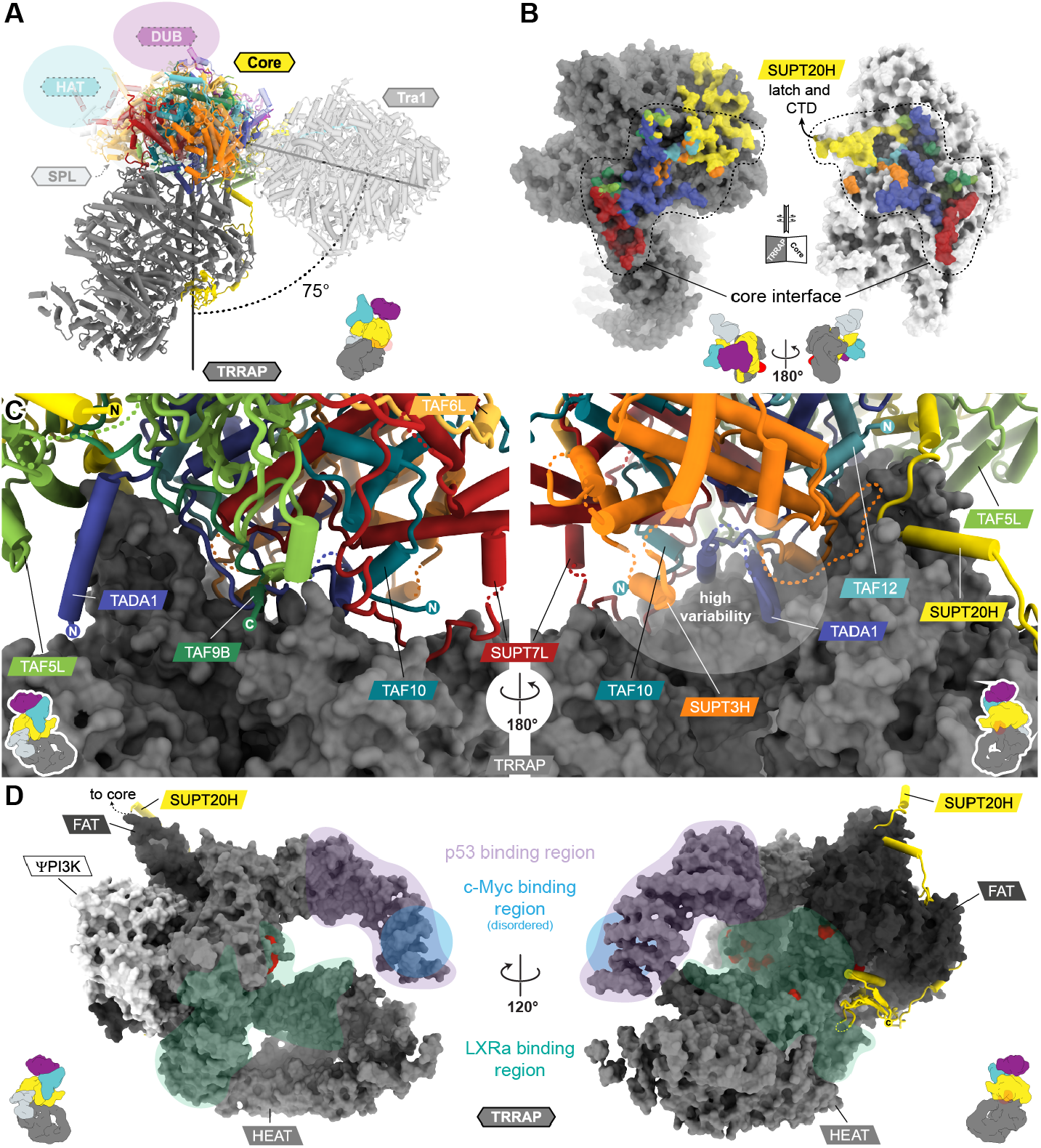
Distinct tethering and interface between TRRAP and the hSAGA core. (**A**) Superposition of the hSAGA (solid) and ySAGA (*15*) (transparent) cryo-EM models on the core module. The approximate location of the remaining modules (transparent circles) is indicated. The angle was calculated between the center of masses of the cores and (pseudo-)kinase domains. (**B**) View on the opened-up TRRAP-core interface of hSAGA (dotted area) in surface representation. Interfacing areas on the TRRAP (left) are colored by the interfacing core subunits they interact with (right). (**C**) Close-up views of the hSAGA TRRAP-core interface from opposite directions. The TRRAP is shown in surface representation, while core subunits are shown in cartoon representation. A region of high variability is indicated by the white oval. (**D**) Activator binding sites mapped on the TRRAP (colored areas) and location of disease mutations (red spots). Schematic insets indicate the viewing direction in all panels.

The best characterized function of SAGA’s TRRAP module, is serving as an interaction hub for transcriptional activators and thus playing a critical role in many cancers and diseases (*10, 30, 41-45*). The SUPT20H CTD resembles a lid that covers a solvent filled tunnel below the FAT domain (Fig. 4C). This tunnel exhibits a strong positive charge (Fig. S10A, B), is highly conserved in metazoans (Fig. S10C, D), and could serve as a binding interface for acidic effector proteins. The metabolite Inositol hexakisphosphate (InsP_6_) was copurified with hSAGA and is bound by highly conserved residues of the FAT and ΨPI3K domains within a side pocket of this tunnel, where it may serve a similar stabilizing role as proposed for mTORC2 (*46*) (Fig. S10E-I).

Structurally, TRRAP appears to display a high degree of conformational flexibility within the HEAT repeat region around the N-terminus and the neighboring cradle. The mapped c-Myc and p53 binding sites on TRRAP (*30, 41*) are located in this flexible region (Fig. 5D, Fig. S11A), which could indicate that c-Myc/p53 binding may stabilize or mediate conformational changes in this part of the structure. Further protein-protein interfaces are found in the FAT proximal HEAT repeat region, where the N-terminal HEAT repeat arm and circular cradle meet. Previous studies identified this region as crucial for interaction with the liver X receptor alpha (LXRa) (*44*) and a cluster of disease-causing missense mutations lie along this region, which is highly conserved in metazoans (*36, 42, 45*) (Fig. 5D, Fig. S11B,C). A number of these mutations, including the well-described recurrent melanoma mutation S722F (corresponding to S721F in the mapped TRRAP isoform), are part of a highly conserved surface patch and likely involved in effector binding (Fig. S11C-E). Other mutations are buried and likely to interfere with the structural integrity of TRRAP (Fig. S11F). Two mutations identified in patients with intellectual disability and neurodevelopmental disorders, map outside of this HEAT repeat cluster (*42*): F859L is located directly at the interface with the newly characterized SUPT20H CTD (Fig. S11G), and R3746Q forms a salt bridge with the highly conserved D291 (Fig. S11H) of the SUPT20H latch on the TRRAP surface (Fig. S7B). Notably, because TRRAP is a scaffold for other important cellular complexes, including the GNC5-related HAT complexes PCAF and ATAC, the MYST family complex TIP60, and the DNA-damage repair MRN complex (*36, 47*), disease-causing mutations may also result in complex-specific perturbations.

### Splicing Module

We could resolve the architecture of the metazoan-specific SPL module within hSAGA from our negative stain studies. The hSAGA subunits SF3B3 and SF3B5 are shared components with the metazoan spliceosome. These proteins are part of the SF3b core complex within the U2 small nuclear ribonucleoprotein (snRNP) involved in branch point sequence recognition during pre-mRNA splicing (*48*). Very little is yet known about how the SPL module functions within SAGA, or how its components partition between SAGA and the U2 snRNP, but more structural relationships between the splicing and transcriptional machinery are emerging (*49*). A recent study also connects ySAGA with splicing through a non-conserved interaction of Spt8 with the spliceosomal ATPase Prp5p (*50*), which could suggest that both complexes have evolved divergent mechanisms to associate with U2 snRNP components and potentially affect co-transcriptional splicing.

The crystal structure of the SF3b complex (*19*) includes three WD40 beta propellers within the large SF3B3 subunit that fit precisely into the EM density, allowing SF3B3 and SF3B5, which bind centrally between those three propellers, to be docked into our hSAGA density map (Fig. 1C, 1D, 3A, S2D, S12A). In the SF3b complex, the two N- and C-terminal propellers of SF3B3, as well as SF3B5, bind to the inside of the circular HEAT repeat (solenoid) subunit SF3B1, which is absent in hSAGA (Fig. S12B). Although using a slightly different interface, within hSAGA this interaction is replaced with the concave face of the TAF6L HEAT repeat. Yeast two-hybrid assays of Drosophila homologs suggested interactions between the SPL, the HAT subunit SGF29, and the core subunit SUPT7L (*51*). Such interactions are consistent with the position of the TAF6L HEAT repeat, which tethers the HAT module (*15, 16*) and directly interacts with the N-terminus of SUPT7L in the core of hSAGA (Fig 3A, 3B, 2A, S6A-C).

The lack of a SPL module in yeast coincides with the presence of short introns and well-defined splice sites, while metazoan introns are long, have poorly conserved splice sites, and are often subject to alternative splicing (*52*). Since splicing occurs primarily co-transcriptionally (*53*), a spatial coordination between splicing, chromatin modification, and transcription has been proposed (*54*). The structure of hSAGA reveals a complex and dynamic architecture, with modules that likely connect these various functions.

Our study shows that hSAGA has a global architecture that is distinct from that of ySAGA. It builds on common elements, such as the distorted HF octamer-like core, but incorporates new ones, such as the SPL module. Unlike the close enhancer-promoter distances found in yeast, metazoan gene regulation contends with a unique transcriptional and chromatin landscape, with kilo- to megabase distances between enhancers and promoters, as well as with promoter architectures and intron and splice site properties that are very distinct from those in yeast (*55-57*). Our hSAGA structure reveals the integration of the splicing module, architectural differences with ySAGA, and distinct tethering of the activator binding subunit that likely reflect unique mechanisms for this complex in human transcription and chromatin regulation.

## Acknowledgments

We thank J. Fang for help with hSAGA purification, A. B. Patel for discussion and the GO protocol, P. Grob and D. Toso, J. Remis, P. Tobias in the Cal-Cryo facility at UC Berkeley for microscope access and support, A. Chintangal for computing support, E. Spooner and the Whitehead MS facility for MS analysis, S. Zheng for help with HeLa cell culture, E. Borbon for help with HeLa cell harvesting, and C. He for help with cloning of constructs and cell maintenance.

## Author contribution

R.K.L piloted cell culture work, purifications, initial negative stain EM, and generated the cell line together with C.D. G.D. cloned CRISPR plasmids and performed genotyping of the edited cell line. M.N.E. performed large scale cell culture and nuclei harvesting, purified hSAGA, performed biochemical validation experiments and analyzed the mass spectrometry data. D.A.H. carried out EM sample preparation, electron microscopy, data processing, model building and refinement, structural analysis, and sequence analysis. Q.F. processed the TBP data set. D.A.H., M.N.E., and E.N. wrote the manuscript.

## Data and materials availability

Cryo-EM maps and refined coordinates were deposited in the Electron Microscopy Data Bank with accession codes EMD-23027 and EMD-23028 and in the Protein Data Bank with accession codes 7KTR and 7KTS. Custom computer code is available from the corresponding author upon reasonable request.

## List of Supplemental Materials

Materials and Methods

Figures S1-S12

Tables S1-S4

Movie S1 References (*58-82*)

