## Supplementary material for "Structure of the human SAGA coactivator complex: The divergent architecture of human SAGA allows modular coordination of transcription activation and co-transcriptional splicing": suplemental figures

**This PDF file includes:**

Figs. S1 to S12
Tables S1 to S4
Caption for Movie S1

**Other Supplementary Materials for this manuscript include the following:**

Movie S1

### Materials and Methods

#### SUPT7L-Halo-(FLAG)3 knock-in cell line generation

Human HeLa cells were cultured at 37 ºC and 5 % $CO_2$ in 4.5 g/L glucose DMEM supplemented with 10 % Fetal Bovine Serum and 10 U/mL Penicillin-Streptomycin and subcultured at a ratio of 1:3 to 1:8 every 2 to 4 days. Genome editing was performed as described previously (*58*).Wild-type HeLa cells were transfected using Lipofectamine 2000 (ThermoFisher Catalog no: 11668019) according to manufacturer's protocol, co-transfecting a Cas9 and a repair plasmid (18 ug repair vector and 6 ug Cas9 vector per P100 dish; 1:3 w/w). The Cas9 plasmid expressed 3xFLAGSV40NLS-pSpCas9-NLS from a CBh promoter; the sgRNA was expressed from a U6 promoter; and mVenus was expressed from a PGK promoter. The repair vector was modified using a pUC57 plasmid to contain the tag of interest (Halo-(FLAG)3) flanked by ~800 bp of genomic homology sequence on either side. Four sgRNAs were designed using the Zhang lab CRISPR design tool (https://zlab.bio/guide-design-resources) cloned, into the Cas9 plasmid, and each sgRNA plasmid was co-transfected with the repair vector individually. After 18–24 h, cells transfected with each of the sgRNAs individually were combined and FACS-sorted for YFP (mVenus) fluorescence, indicating successful transfection. YFP-sorted cells were grown for 4–12 days, labeled with 500 nM Halo-TMR and cell populations with higher fluorescence than similarly labeled wild-type cells were FACS-selected and sorted individually into 96-well plates. Clones were expanded and genotyped by PCR using a three-primer PCR (genomic primers external to the homology sequence and an internal Halo primer). Successfully edited clones were further verified by PCR with multiple primer combinations, Sanger sequencing and Western blotting.

#### Preparative HeLa cell culture and nuclei extraction

Large scale cultures of SUPT7L-Halo-(FLAG)3 HeLa cells were grown at 37 ºC and ambient $CO_2$ in a Hotpack Environmental Chamber (Scientific Products) in Joklik-modified Minimum Essential Medium Eagle (Sigma) media supplemented with 5 % Bovine Calf Serum, 50 U Penicillin-Streptomycin, and 2 mM Glutamax (Thermo Fisher). Cells were maintained in 6 L Florence round-bottom spinning flasks (Fisher Scientific) each containing 4 L of HeLa cultures and constantly stirred via a Precision Magnetic Stirrer Platform (Belloco). Every 24 h, cells are split 1:2 into fresh media grown to a density of ~$2.5\text{-}5 \times 10^5$ cells/mL, and harvested (up to 16 L). To harvest, SUPT7L-Halo-(FLAG)3 HeLa cells were centrifuged using a Fiberlite F9-6 x 1000 LEX Fixed Angle Rotor (Thermo Fisher) at 4 ºC and 4,000 rpm for 15 min. Cells were washed in PBSM (PBS buffer with 5 mM $MgCl_2$) then centrifuged using a Eppendorf A-4-62 Swinging Bucket Rotor at 3,800 rpm for 10 min. Cells were resuspended in 5 volumes of Buffer A (10 mM HEPES pH 7.6, 1.5 mM $MgCl_2$, 10 mM KCl, 1x Roche cOmplete protease inhibitors) briefly vortexed, incubated on ice for 20 min, and centrifuged (Eppendorf A-4-62, 3,800 rpm,10 min, 4 ºC). Cells were lyzed by resuspension in 2 volumes of Buffer A and douncing with 7 strokes using a glass homogenizer with a Type "B" pestle. Nuclei were pelleted by centrifugation (Eppendorf A-4-62, 2,700 rpm, 10 min, 4 ºC), flash frozen in liquid nitrogen, and stored at -80 ºC until use.

#### hSAGA purification

All steps were performed at 4 ºC. Frozen nuclei from ~30-40 L cell culture were thawed, 0.9 volumes of Buffer C (20 mM HEPES pH 7.8, 1.5 mM $MgCl_2$, 0.2 mM EDTA, 25 % glycerol, 0.42 M KCl, 1 mM DTT, 0.5 mM phenylmethylsulfonyl fluoride (PMSF) and 1 uM Leupeptin) added, and dounced using a glass homogenizer and a type "B" pestle 20 times on ice. The nuclear extract was then centrifuged using a JA-20 Beckman rotor at 4 ºC and 20,000 rpm for 30 min. The supernatant was collected and the conductivity adjusted to the conductivity of 0.3 M NaCl at 4 °C. The nuclear extract (~60 ml) was loaded on a pre-equilibrated (0.3 M NaCl HEMG (20 mM Hepes-KOH pH 7.6, 2 mM $MgCl_2$, 0.2 mM EDTA, 20% glycerol) buffer) 50 mL phosphocellulose P11 (GE Healthcare/Whatman) column. Unbound protein was eluted using 3 CV of 0.3 M NaCl HEMG. Bound protein was eluted in two steps with 3 CVs 0.5 M NaCl HEMG, followed by 3 CVs of 1.0 M NaCl HEMG, and fractionated (5 mL). Peak fractions were determined by Bradford assay and combined. Human SAGA eluted with the 0.5 M NaCl HEMG peak (hereafter called P0.5M) and dialyzed against 150 mM KCl Buffer D (20 mM HEPES pH 7.8, 2 mM $MgCl_2$, 0.2 mM EDTA, 10 % glycerol, 150 mM KCl, 0.5 mM PMSF and 1 uM Leupeptin) using SnakeSkin 10 kDa MWCO dialysis tubing (Thermo Fisher). The dialyzed P0.5M fraction was supplemented with IGEPAL CA-630 (0.1% (v/v) final) and incubated with 500 uL beads of pre-equilibrated FLAG M2 resin (Sigma) for 12 h on a nutating platform. The resin was washed twice with 2 CV of Column Buffer (25 mM HEPES pH 7.8, 0.2 M NaCl, 10% (v/v) glycerol, 1 mM EDTA, 0.5 mM TCEP, 0.1% (v/v) IGEPAL CA-630, 1x Roche cOmplete protease inhibitors ), twice with 2 CV Wash Buffer (25 mM HEPES pH 7.8, 0.6 M NaCl, 10% (v/v) glycerol, 1 mM EDTA, 0.5 mM TCEP, 0.1 %(v/v) IGEPAL CA-630, 1x Roche cOmplete protease inhibitors), and twice with 2 CV Column Buffer. Protein was eluted from the FLAG resin by adding Column Buffer with 0.1 mg/mL 3xFLAG peptide and incubating on a rocking platform for 1 h. The hSAGA containing supernatant was removed after centrifugation (Eppendorf 022653041 fixed-angle rotor, 3,300 rpm, 5 min). The elution was repeated four times. The first 3 elutions were concentrated 5-fold using a 100 kDa MWCO Spin-X UF concentrator (Corning) and used for biochemical and EM studies. The sample was frozen in liquid nitrogen and stored at -80 ºC. Sample quality and the effect of freeze-thaw cycles was analyzed by negative stain EM throughout the purification. All FLAG elution fractions yielded similar quality in cryo-EM. A concentration of approx. 50 nM was determined by densitometry.

**Mass Spectrometry**

Samples of purified hSAGA were shipped overnight on dry ice from Berkeley, CA and prepared and analyzed by mass spectrometry by the Whitehead Institute Proteomics Facility (Cambridge, MA). Samples were diluted to 100 µL in 6 M Urea, 100 mM Tris pH 7.8 buffer. Dithiothreitol (DTT, 5 µL of 200 mM) was added and incubated for 60 min at room temperature. Free cysteines were alkylated by addition of 20 µL of 200 mM iodoacetamide and incubated for 60 min at room temperature. The urea concentration was lowered by adjusting the sample volume to 900 µL with 100 mM Tris pH 7.8 buffer. The protein was digested by adding 100 µL of a 20 ng/uL trypsin or chymotrypsinin solution and incubated overnight at 37 °C with gentle shaking. The resulting peptides were washed, extracted and concentrated by solid phase extraction using Waters Sep-Pak Plus C18 cartridges. Organic solvent was removed and volumes reduced to 15 µL for subsequent analyses using a SpeedVac operated at 60 °C. The digested extracts were analyzed by reversed phase high performance liquid chromatography (HPLC) using Waters NanoAcquity pumps and autosampler and a ThermoFisher Orbitrap Elite mass spectrometer using a nano flow configuration operated in a data dependent manner for the 60 min. The resulting fragmentation spectra were correlated against the Uniprot isoforms and TrEMBL databases for *homo sapiens* using Sequest (Thermo Fisher Scientific, San Jose, CA, USA; version IseNode in Proteome Discoverer 1.4.1.14).

Sequest was searched with a fragment ion mass tolerance of 0.50 Da and a parent ion tolerance of 15 PPM. Carbamidomethyl of cysteine was specified in Sequest as a fixed modification and oxidation of methionine was specified as a variable modification. Scaffold (version Scaffold_4.11.0, Proteome Software Inc.) was used to provide consensus reports for the identified proteins. Peptide identifications were accepted if they could be established at greater than 95.0 % probability by the Scaffold Local FDR algorithm. Protein identifications were accepted if they could be established at greater than 99.0 % probability and contained at least one identified peptide.

**Negative stain sample preparation of hSAGA, data collection, and processing**
400 mesh Cu grids were cleaned three times (in ethanol, water, ethanol) by sonication for five min and dried on filter paper. A petri was filled with ultrapure water forming a meniscus and wiped off with lens paper. One drop of 1 % (w/v) nitrocellulose in amyl acetate was added to the surface, forming a thin film. Cleaned grids were deposited on the film with the polished side facing down. The grids were transferred with parafilm onto filter paper with the nitrocellulose facing up and dried overnight. Grids were coated with carbon by evaporation using an Edwards Auto306 ($10^{-6}$ mbar, 6 A, 6 sec). Prior to sample adsorption, grids were glow discharged (30 s, 15 W, Tergo EM PIE scientific). Human SAGA was diluted (2x) in dilution buffer (25 mM HEPES pH 7.5, 0.2 mM EDTA, 6 mM $MgCl_2$, 0.2 M NaCl, 3 % (w/v) D(+) Trehalose), 3 µl were applied to a grid and adsorbed for one min. The grid was washed and stained, respectively, by swirling it five times on a 50 µl drop of 2 % (w/v) uranyl formate for 10 sec (each), blotted, and dried in an air stream. Data was collected on a Tecnai F20 (Thermo Fisher Scientific), operated at 120 kV, and equipped with an UltraScan4000 (Gatan) using Leginon (*59*) (Table S4, Fig. S2), with a pixel size of 1.4 Å, using a defocus range of 0.4-3.9 µm, and a total dose of 35 $e^{-}Å^{-2}$. Micrographs were contrast transfer function (CTF) corrected using CTFfind 4.1.13 (*60*). Particles were picked using gaussian LoG picker in Relion-3.1 (*61*), extracted with a box size of 300x300 pixels, and subjected to reference free 2D classification. Particles from the best classes (32 %) were used for initial model generation using the statistical gradient descent method (*62*) in Relion-3.1 (*61*). Particles were classified by a series of 3D and 2D classifications with and without alignment (Fig. S2A). Classification revealed one class without (ordered) TAF6L HEAT domain and SPL module. Particles with and without this region were separated by multi-reference 3D classification. The best reconstruction was refined and classified again by alignment free 3D classification. Combined classes that yielded the highest resolution were refined, and postprocessed in Relion-3.1 (*61*) using a sharpening B-factor of -1,200 $Å^2$ (Fig. S2B).

**Cryo-EM sample preparation of hSAGA, data collection, and processing**
Quantifoil Au 300 mesh UltrAuFoil R1.2/1.3 polyethylenimine (PEI) / graphene oxide (GO) grids were prepared using the following procedure: Grids were washed twice with chloroform, glow discharged on glass (Tergo EM PIE scientific, 15 W, 59 sec), and coated on non-capillary tweezers with polyethylenimine (PEI) by incubation with 4 µl of freshly prepared 1 mg/ml PEI (MAX linear MW 40k) in 25 mM HEPES pH 7.9 for two min. The PEI solution was blotted off, followed by washing and blotting twice with 4 µl of water without letting the grid dry. The grids were then dried for two min on tweezers and 15 min on filter paper (PEI facing up). PEI coated grids were incubated on non-capillary tweezers with 4 µl of freshly prepared 0.2 mg/ml GO suspension (1:10 dilution of Sigma 763705, centrifuged at 1,500 g) for 2 min, blotted with a linear motion along the

blotting paper, washed twice (see above), and dried for one min on tweezers. The grids were used for freezing within 2-4 h.

All grid preparation steps were done on ice. 3 µl of undiluted hSAGA was transferred into a 0.5 ml non-stick tube and crosslinked by mixing with 0.6 µl of crosslinking buffer (25 mM HEPES pH 7.8, 0.2 M NaCl, 0.2 mM EDTA, 0.5 mM TCEP, 0.01 % (v/v) NP40, 10 % (v/v) glycerol, 6 mM bis(sulfosuccinimidyl)suberate (BS3)), and incubated for five min. A GO grid was picked up with Vitrobot tweezers, the sample was transferred to the grid and incubated for two min in a saturated humidity chamber. Afterwards, the grid was washed five times by submerging and swirling for 5 sec (each) in 230 µl of wash buffer (25 mM HEPES pH7.8, 0.2 M NaCl, 0.2 mM EDTA, 0.5 mM TCEP, 0.01 % (v/v) NP40, 2.5 % (v/v) glycerol) in a five well Teflon block. Without letting the grid dry, excess solution was blotted off at a 90° angle and 4 µl wash buffer were added immediately. The grid and tweezers were mounted into a Vitrobot Mark IV (Thermo Fisher Scientific), blotted with fresh filter paper (blot force 0, 3 sec), and plunge frozen into liquid ethane.

Data was collected with SerialEM (*63*), with a physical pixel size of 1.187 Å, a defocus range of 0.9-3.4 µm, a total dose of 50 $e^{-}Å^{-2}$, and 3x3 multishot acquisition on a Titan Krios G2 (Thermo Fisher Scientific), operated at 300 kV, with BIO Quantum Energy Filter (Gatan) and a K3 Direct Electron Detector (Gatan) operated in super resolution mode (Table S4). Movies were whole-frame motion corrected and binned (2x) using the Relion-3.1 (*61*), CTF corrected using CTFfind 4.1.13 (*60*), and sorted manually. Particles were picked using the gaussian LoG picker in Relion-3.1 (*61*) and extracted with 8x binning (Fig. S3A) and a box size of 45x45 pixels. GO edges were removed by 2D classification before hSAGA particles could be classified. After removing the majority of GO edges, particles were re-extracted with recentering (4x binned, 90x90 box size) and re-classified in 2D. The negative stain reconstruction was low pass filtered to 50 Å and used as reference model for initial 3D classification. Each class was sub-classified by alignment free 2D classification to remove particles close or on GO edges. The remaining particles were subjected to 3D classification, re-centered in the box by applying a coordinate transformation to the particle alignment parameters using python, re-extracted with re-centering, without binning, and a box size of 360x360 pixels, and subjected to a consensus 3D refinement. Afterwards, the particles were subjected to two rounds of Bayesian polishing, 3D refinement, CTF refinement, and alignment free 3D classification (tau=20) (Fig. S3A). A final round of 3D classification, refinement, and postprocessing yielded a reconstruction at 2.93 Å (Fig. S3B-D). High variability and low local resolution were observed at the TRRAP N-terminus and the HEAT repeat cradle in close proximity as well as around the surface of the core module. Low pass filtering and B-factor blurring slightly improved the interpretability of the map in these regions. Further improvement was made by multibody refinement (Fig. S3E, S4D) of the core and TRRAP modules, although the resolution did not improve. Various density modification and map enhancement methods were tested, and the most significant improvement in variable and surface exposed regions were obtained by applying the spiral phase transform in LocSpiral (*64*) to all reconstructions. This process revealed additional peptide connections and a density fragments of the disordered TAF6L HEAT domain (Fig. S3E, S4A). A principal component analysis of the multibody refinement showed a high degree of flexibility between the core and TRRAP modules (Fig. S4A), which can also be observed by 3D variability analysis in Cryosparc 2.15.0 (*62, 65*) (data not shown). Masking and map transformations were carried out using UCSF Chimera (*66*) and Relion-3.1 (*61*). All resolution estimates are based on the 0.143 threshold criterion of the gold-standard Fourier Shell Correlation

(FSC) (*67*) of two independently refined half sets in Relion-3.1 (*61*), after accounting for correlations introduced by masking (*68*). Local resolution was estimated using Relion-3.1 (*61*).

**Cryo-EM sample preparation of hSAGA with TBP, data collection, and processing**
Human SAGA was mixed with a sixfold molar excess of human full length TBP and incubated for 5.2 h on ice. Grids were frozen and data was collected and processed in the same way as described above, but no additional density corresponding to TBP could be observed.

**Modeling and refinement**
For model building in Coot (*69*) maps were converted to structure factors using phenix.map_to_structure_factors (*70*), allowing low-pass filtering and variable B-sharpening/blurring in Coot (*69*). Models were built into the postprocessed, multibody refined, and LocSpiral (*64*) filtered maps (Fig. S3E, F). A fragmented initial model of secondary structure elements in TRRAP was generated using phenix.map_to_model (*70*), manually corrected, and completed in Coot (*69*). A homology model of the TAF5L WD40 propeller was generated using SwissModel (*71*) (based on PDB 6F3T (*72*)) and rigid body fitted in Coot (*69*). The remaining model was build de-novo in Coot (*69*), guided by homology models based on hTFIID (*5*) and ySAGA (*15, 16*). Regions with low confidence in register assignment were modeled as poly-alanines (assigned as UNKs). Prior to real-space refinement in Phenix, atomic B-factors were reset to 90 Å$^2$, the model was protonated using phenix.ready_set (*70*), and sanity checked as well as geometry minimized using gelly (*73*) (GlobalPhasing). Afterwards, the model was refined using Rosetta (*74*), validated using phenix.molprobity (*70*), and optimized in Coot (*69*). Secondary structure restraints were generated using phenix.secondary_structure_restraints and corrected after manual inspection. A final refinement was carried out with phenix.real_space_refine (*70*) (1.18-3861) against the complete LocSpiral (*64*) filtered map using default parameters plus secondary structure restraints, rotamers.fit=outliers_and_poormap, and rotamers.tuneup=outliers_and_poormap (Fig. S3F). Model statistics were calculated using phenix.molprobity (*70*) (Table S4). Refinement against the regular postprocessed map resulted in almost identical statistics with an all-atom r.m.s.d. of 0.400 Å. All maps used for model building and refinement were deposited in the EMDB. Map vs. model Fourier shell correlation (FSC) was calculated using phenix.mtriage (*70*) (Fig. S3G). The $InsP_6$ ligand was identified by density fit and homology to mTORC2 (*46*) and one out of two possible conformations was modeled (Fig. S10E). In analysis of our cryo-EM structure, separating the core and TRRAP modules improved map quality and revealed additional features on surface exposed regions after LocSpiral filtering (*64*) (Fig. S4A). In particular, analysis of the region where SUPT3H, SUPT7L, TADA1, and TRRAP meet suggested alternative main chain conformations that could not be sorted out by classification. Importantly, the highly variable region between SUPT3H, TADA1, and TRRAP corresponds to the approximate position where TBP binds to ySAGA (Fig. S4B,C).
A model for the negative stain reconstruction was generated by rigid body fitting without coordinate refinement in phenix.real_space_refine (*70*) using the protein part of the cryo-EM model, a homology model of the TAF6L HEAT domain (generated with SwissModel (*71*) and based on human TAF6 (*5*), PDB 6MZL), and SF3B3/SF3B5 from the SF3b core complex (*19*) (PDB 5IFE) (Fig. S2D, Table S-A). Prior to fitting, expression tags in SF3B3 were deleted and the TAF6L HEAT domain was mutated to poly-alanines (annotated as UNKs) to reflect the absence of an authentic high- or medium-resolution structure for this region.

**Structural analysis and visualization**

Coordinate transformations and manipulations were carried out using CCP4 tools (*75*). Structures were compared using PDBefold (*76*) and interfaces were analyzed using QtPISA v2.1.0 (*75*). Relative angles between variable regions/domains (*e.g.* TAF5(L) NTDs) of related structures with a common reference domain (*e.g.* TAF5L WD40) were calculated by pre-aligning all structures to the reference domain of hSAGA using secondary structure matching (SSM). The center of masses of the hSAGA reference domain (*e.g.* TAF5L WD40), the hSAGA variable domain (*e.g.* TAF5L NTD), and the hSAGA variable domain after superposition on the corresponding domain in related structures using SSM (*e.g.* ySAGA NTD) were calculated. Center of masses were calculated in PyMOL (The PyMOL Molecular Graphics System, Version 2.4.0 Schrödinger, LLC.) and angles between corresponding vectors were calculated using python. Structure figures were generated using PyMOL, ChimeraX (UCSF, 2020-01-10), and Adobe Illustrator. Electrostatic surfaces were generated using the APBS (*77*) plugin in PyMOL. Videos were generated using ChimeraX (*78*) (UCSF, 2020-01-10), Adobe Premier, and ffmpeg (https://ffmpeg.org). Plots were generated using python. Reported contour levels for maps are defined as $\sigma$ = density threshold / r.m.s.

**Sequence analysis**

23 metazoan homologs of hSAGA with a complete set of all 20 subunits were retrieved from databases. Sequence alignments were generated using the Clustal Omega (*79*) executable in Geneious Prime 2021.0.3. Phylogenetic trees were generated using the neighboring joining algorithm in Geneious Prime 2021.0.3. Sequence conservation figures were generated by aligning all subunit sequences of all 23 metazoan SAGAs with the sequences of the molecular model of hSAGA. Alignments were combined and conservation scores were calculated using AL2CO (*80*) and used for coloring in PyMOL.

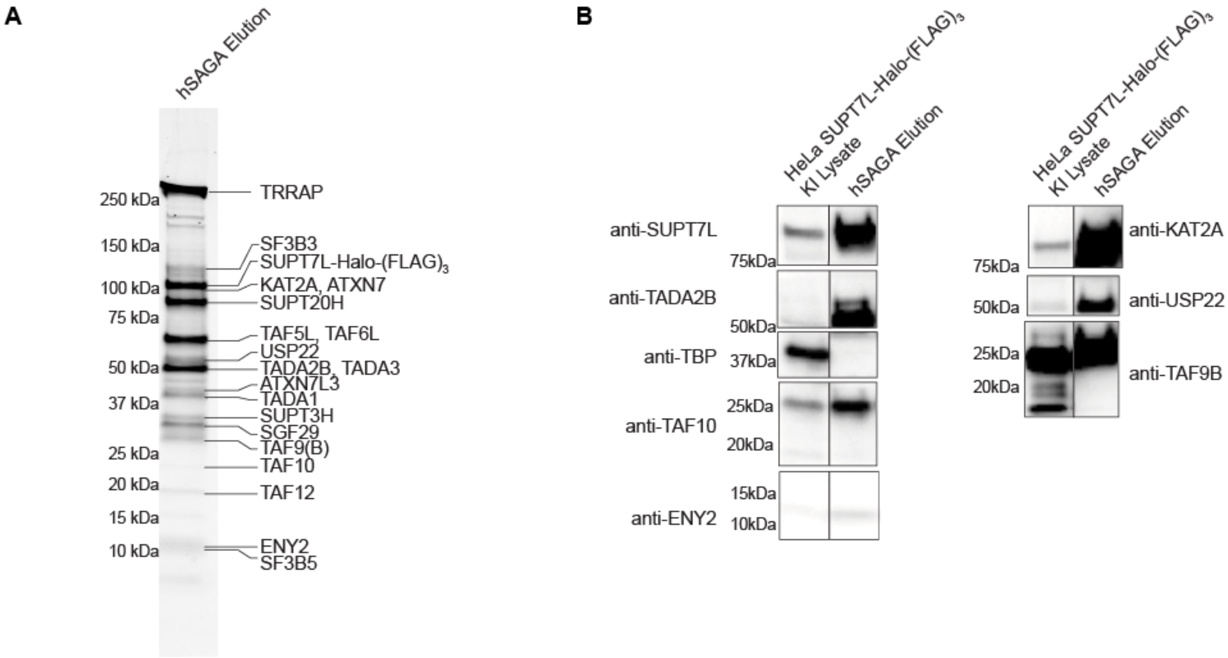

**Fig. S1. Purification of hSAGA.** (**A**) hSAGA elution with subunits labeled based on their predicted molecular weight. (**B**) Western blot probing for DUB, HAT, and core subunits to verify the presence of these modules in the sample used for grid preparation. TBP did not significantly co-purify with hSAGA. KI: knock-in. Blots were cropped.

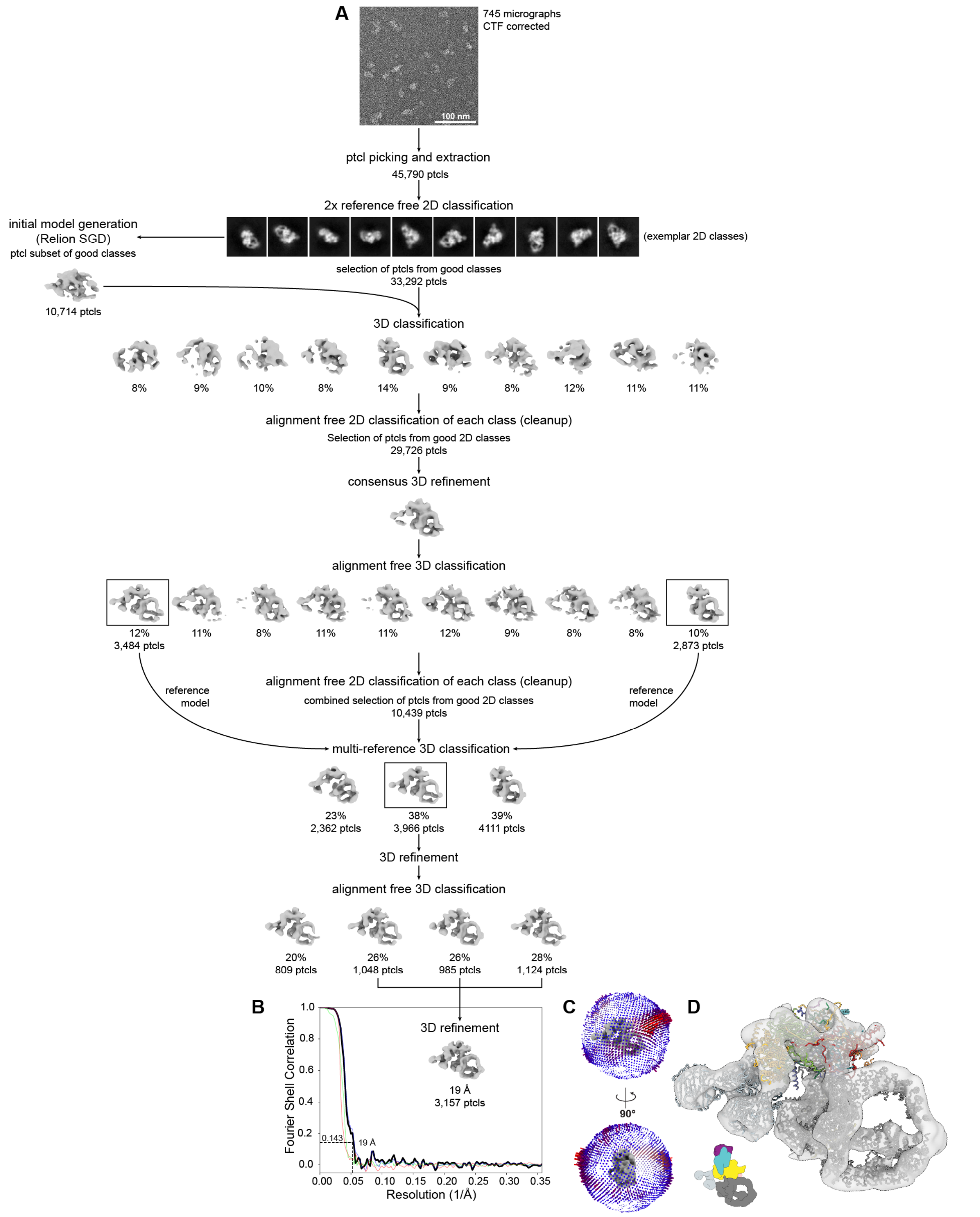

**Fig. S2. Negative stain processing scheme and model fit.** (**A**) After initial 2D classification, particles from the best classes were used for initial model generation in Relion-3.1 (*61*)

implementation of the Cryosparc (*62*) patented stochastic gradient descent (SGD) method. The data was cleaned up by 3D classification followed by alignment-free 2D classification. Particles from all good classes were subjected to a consensus 3D refinement followed by alignment-free 3D classification. All except for one class revealed fuzzy density for the TAF6L HEAT and SPL region. Subsequently, all classes were cleaned up individually by alignment-free 2D classification and combined in a multi-reference classification using the two best models with and without TAF6L HEAT and SPL region. The best class including this region was subjected to 3D refinement, alignment-free 3D classification, and to a final refinement using particles of the class combination that yielded highest resolution. (**B**) Final map and FSC plot. (**C**) Angular distribution. (**D**) Final map (contoured at 4.9 $\sigma$) and rigid body fit of SF3B3/SF3B5 (from PDB 5IFE), a homology model of the TAF6L HEAT domain, and the cryo-EM structure from this study. A schematic colored by modules (see Fig. 1A) in the lower left corner represents the view of all reconstructions in the figure.

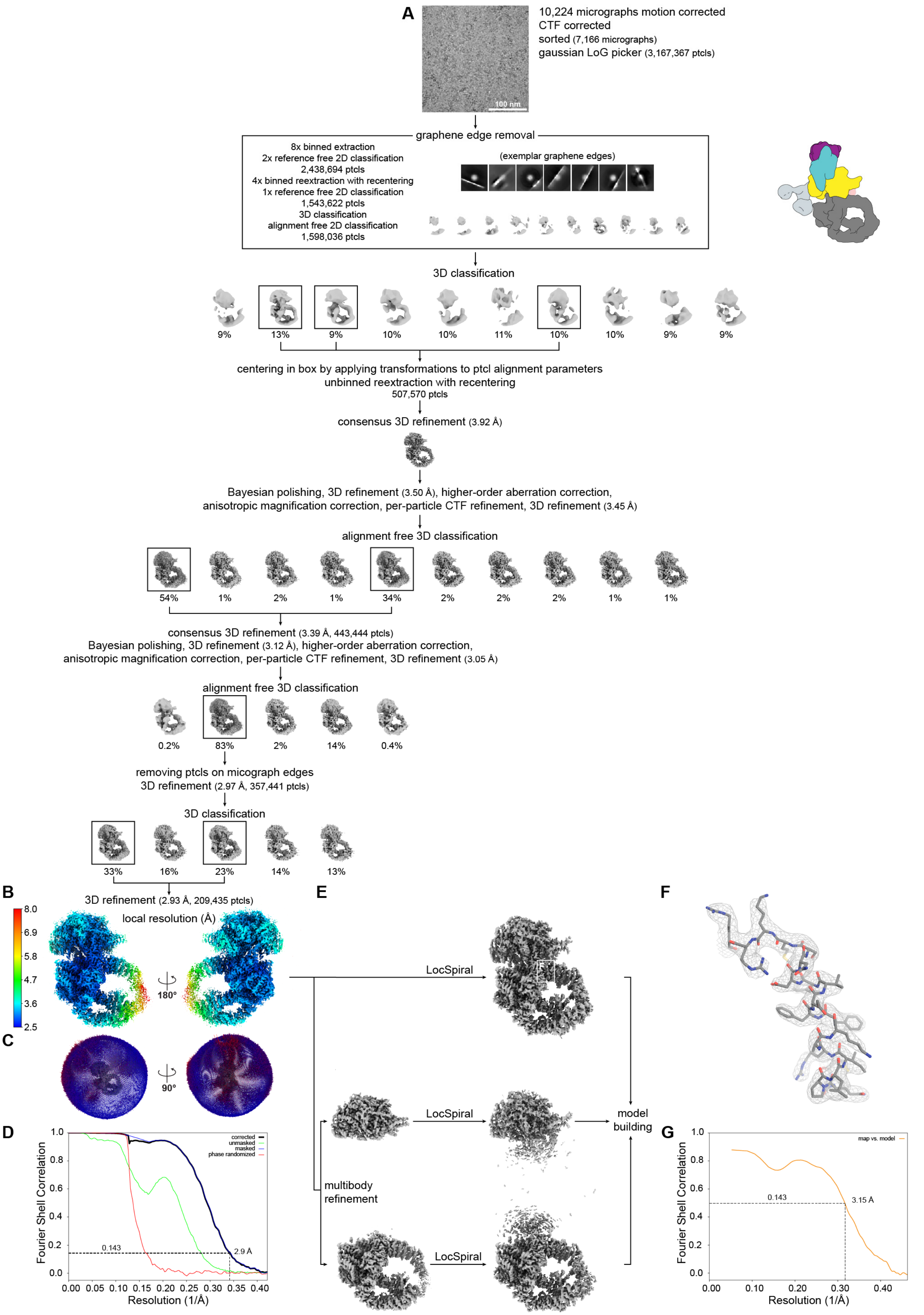

A
10,224 micrographs motion corrected
CTF corrected
sorted (7,166 micrographs)
gaussian LoG picker (3,167,367 ptcls)
100 nm
graphene edge removal
8x binned extraction
2x reference free 2D classification
2,438,694 ptcls
4x binned reextraction with recentering
1x reference free 2D classification
1,543,622 ptcls
3D classification
alignment free 2D classification
1,598,036 ptcls
(exemplar graphene edges)
3D classification
9% 13% 9% 10% 10% 11% 10% 10% 9% 9%
centering in box by applying transformations to ptcl alignment parameters
unbinned reextraction with recentering
507,570 ptcls
consensus 3D refinement (3.92 Å)
Bayesian polishing, 3D refinement (3.50 Å), higher-order aberration correction,
anisotropic magnification correction, per-particle CTF refinement, 3D refinement (3.45 Å)
alignment free 3D classification
54% 1% 2% 1% 34% 2% 2% 2% 1% 1%
consensus 3D refinement (3.39 Å, 443,444 ptcls)
Bayesian polishing, 3D refinement (3.12 Å), higher-order aberration correction,
anisotropic magnification correction, per-particle CTF refinement, 3D refinement (3.05 Å)
alignment free 3D classification
0.2% 83% 2% 14% 0.4%
removing ptcls on micograph edges
3D refinement (2.97 Å, 357,441 ptcls)
3D classification
33% 16% 23% 14% 13%
B
3D refinement (2.93 Å, 209,435 ptcls)
local resolution (Å)
8.0
6.9
5.8
4.7
3.6
2.5
180°
C
90°
D
Fourier Shell Correlation
Resolution (1/Å)
corrected
unmasked
masked
phase randomized
0.143
2.9 Å
E
LocSpiral
LocSpiral
LocSpiral
multibody
refinement
model
building
F
G
Fourier Shell Correlation
Resolution (1/Å)
map vs. model
0.143
3.15 Å

**Fig. S3. Cryo-EM processing and model building scheme.** (**A**) Classification of hSAGA is dominated by the presence of graphene oxide (GO) edges that had to be removed in cycles of initial 2D and one 3D classification. The negative stain reconstruction (Fig. S2) was used as initial model. The relative orientation of all reconstructions in the figure is indicated by the schematic on the top right (see Fig. 1A). 3D classes were centered in the box by applying a coordinate transformation to the alignment parameters, and unbinned particles were re-extracted with re-centering. Particles were filtered for high resolution features in cycles of 3D refinement, classification, Bayesian polishing, and CTF refinement as indicated. Postprocessed map (B-sharpened with -51.1 Å$^2$, contoured at 4.9 σ) with local resolution (**B**), angular distribution (**C**), and Fourier Shell Correlation (FSC) plots (**D**) are shown for the highest resolution class. (**E**) Multibody refinement improved map quality, but not the overall resolution. Considerable improvement of map quality, particularly in surface exposed regions, was achieved by filtering with LocSpiral (*64*). The model for the core and TRRAP was built into the LocSpiral filtered maps of the multibody refinement. The interface between these regions was build using the full map and used for model refinement. Refinement against the postprocessed map (**B**) resulted in the same model, with virtually identical statistics and an all-atom r.m.s.d. of 0.400 Å. Maps are contoured at (regular/LocSpiral): Core 11.2 σ/9.2 σ, TRRAP 7.9 σ/9.0 σ, full 6.9 σ. (**F**) The refined map shows well defined secondary structure elements and side chains (contoured at 9.0 σ). (**G**) Map vs. model FSC using the postprocessed map shown in **B**.

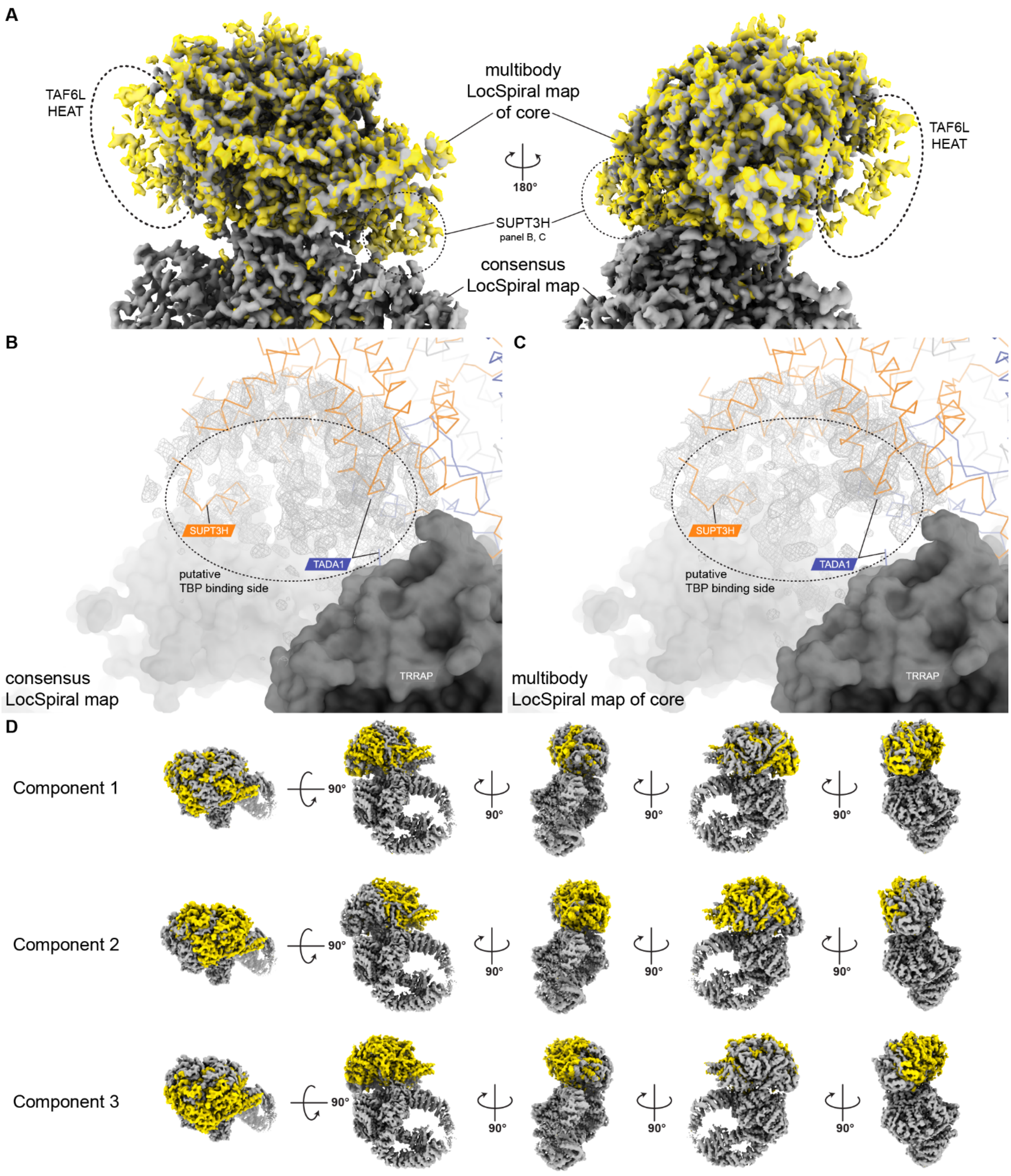

**Fig. S4. Conformational variability in hSAGA between the core and TRRAP domains.** (A) The LocSpiral filtered multibody map of the core reveals additional density corresponding to the poorly ordered TAF6L HEAT domain, and to SUPT3H in the cleft between the core and TRRAP module. All maps are contoured at 5.9 σ. (**B, C**) The cleft between SUPT3H, TADA1 (colored ribbon representation), and TRRAP (surface representation) reveals highly variable density, shown in grey mesh with a radius of 20 Å, in the multibody (B, contoured at 9.2 σ) and the LocSpiral filtered consensus maps of the core (C, contoured at 6 σ). Other core subunits are indicated in light grey ribbon. (**D**) Principle component analysis of the multibody refinement reveals several tilting and swiveling motions between the core and TRRAP module.

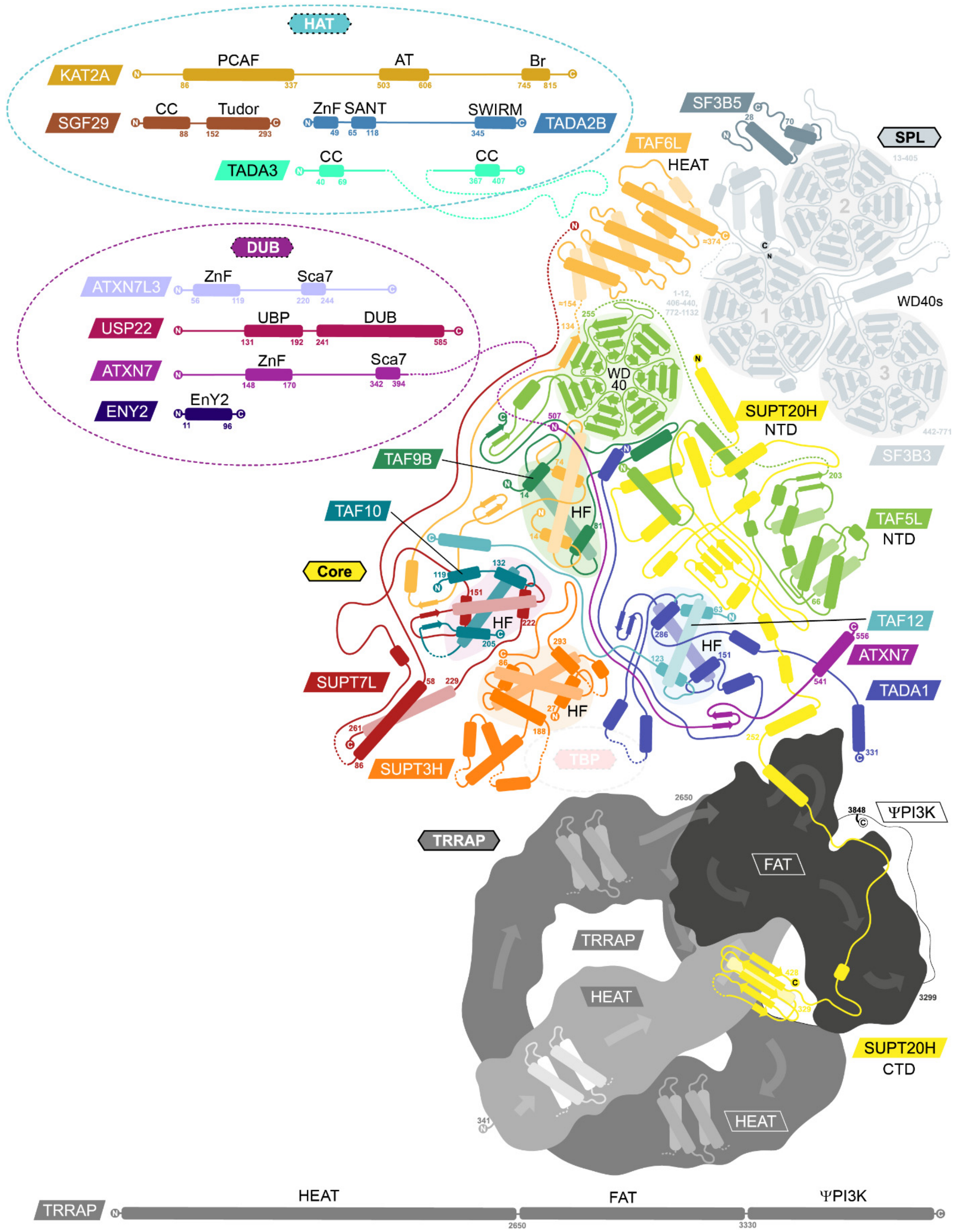

**Fig. S5. Full topology plot of hSAGA.** Schematic showing the relative positions and interactions of hSAGA subunits. WD40 propeller and Histone folds (HF) are indicated by a colored background. Domain boundaries are indicated as residue numbers corresponding to the isoforms described in this paper (see Table S2). No structure of the human HAT or DUB module is available, so domain boundaries for the DUB module are derived from prediction (*81*) and from the HAT module from a published crosslinking mass spectrometry modeling study(*82*). Subunits and domains are not drawn to scale.

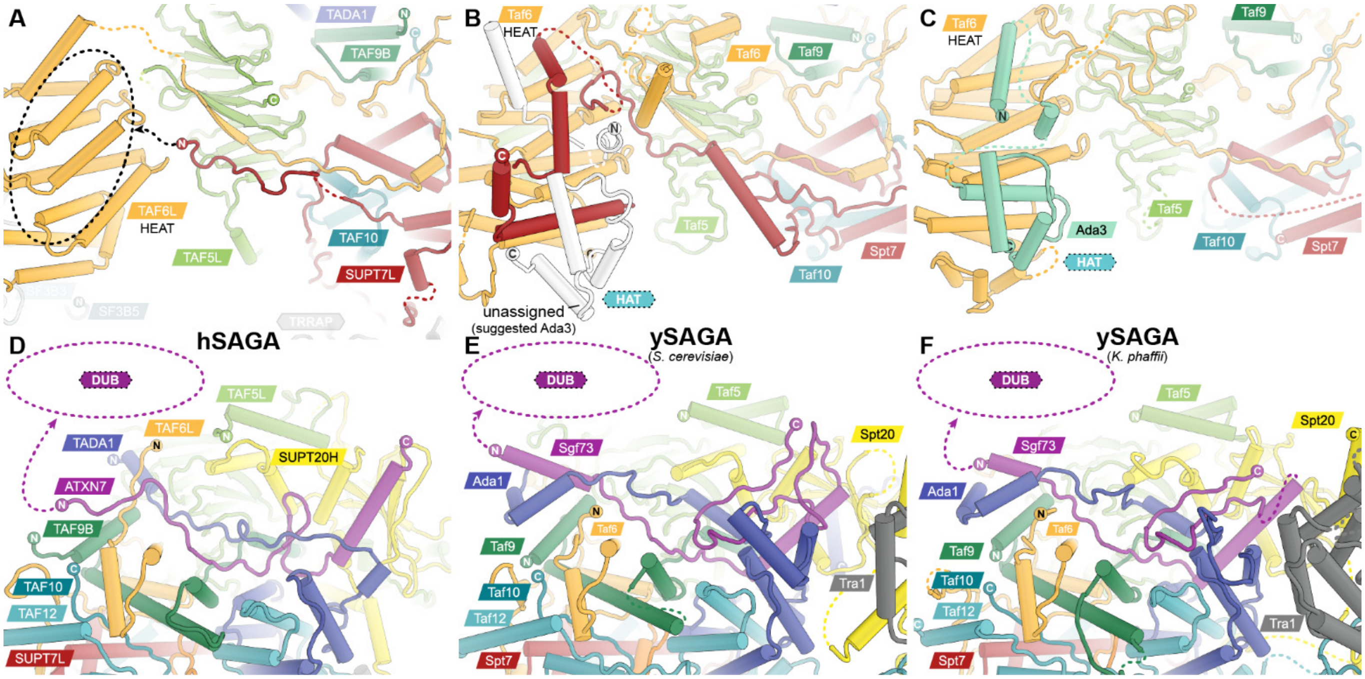

**Fig. S6. The HAT and DUB modules are likely to be positioned in hSAGA similarly to their counterparts in ySAGA.** (**A**) The N-terminus of SUPT7L runs parallel to the TAF6L linker, which connects to the HEAT domain, along the surface of the core and ends with its N-terminus in close proximity to the HEAT domain. (**B**) In the yeast *Saccharomyces cerevisiae* (PDB 6T9I (*15*)), the Spt7 linker further extends towards the convex surface of the TAF6L HEAT domain and interacts with an unassigned region, which was annotated as the Ada3 subunit of the HAT module in (*16*) (**C**). The same regions were unassigned in PDB 6TB4 (*Komagataella phaffii)*. The similar location of the SUPT7L N-terminus suggests a similar interaction and connectivity for the HAT module in hSAGA. (**D**) The ATXN7 subunit of the core and the DUB module is similarly integrated into the core module as in ySAGA (**E**, **F**), suggesting a similar relative attachment of the human DUB.

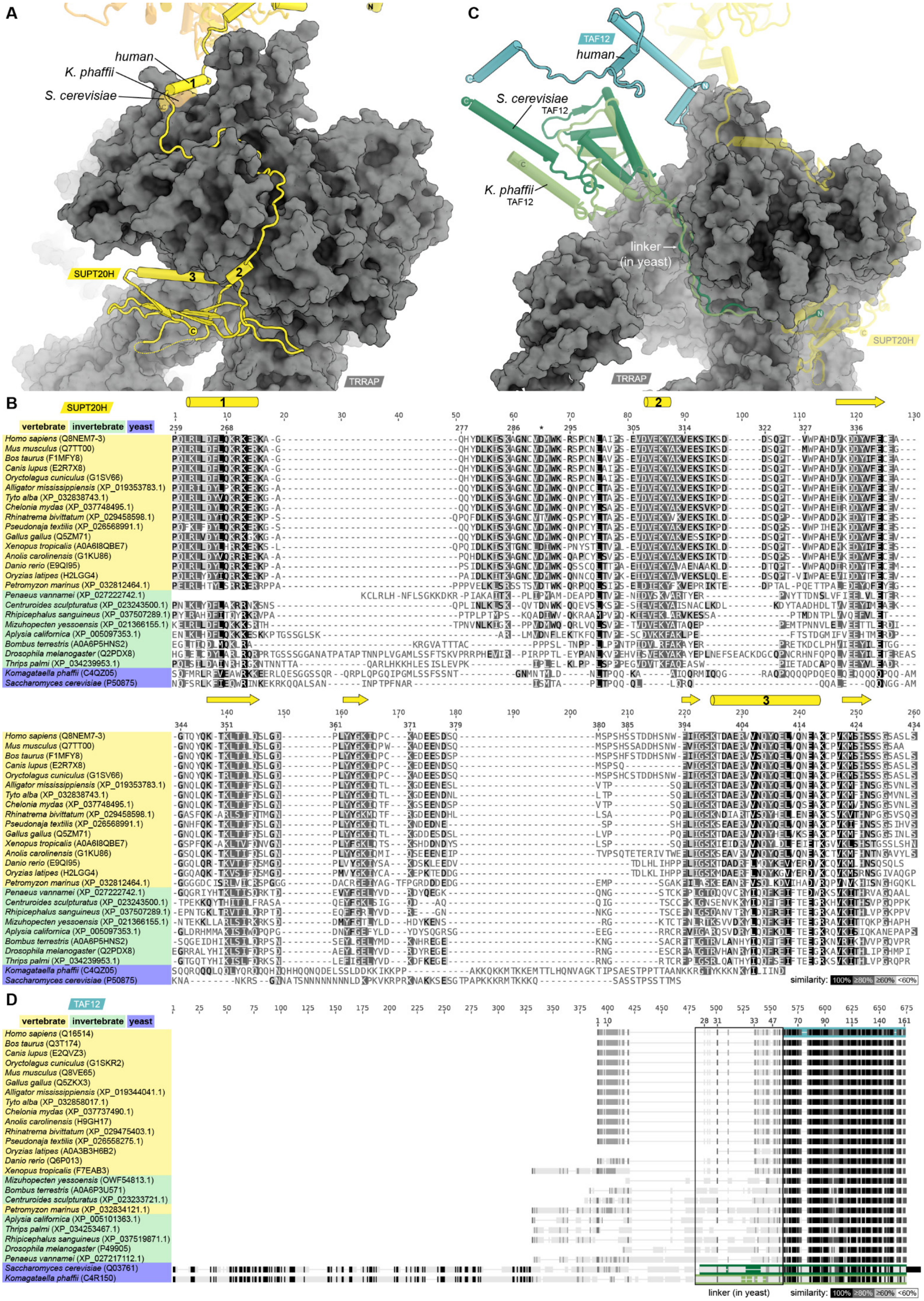

A
human
K. phaffii
S. cerevisiae
SUPT20H
TRRAP
C
TAF12
human
S. cerevisiae TAF12
K. phaffii TAF12
linker (in yeast)
SUPT20H
TRRAP
B
SUPT20H
vertebrate
invertebrate
yeast
Homo sapiens (Q8NEM7-3)
Mus musculus (Q7TT00)
Bos taurus (F1MFY8)
Canis lupus (E2R7X8)
Oryctolagus cuniculus (G1SV66)
Alligator mississippiensis (XP_019353783.1)
Tyto alba (XP_032838743.1)
Chelonia mydas (XP_037748495.1)
Rhinatrema bivittatum (XP_029458598.1)
Pseudonaja textilis (XP_026568991.1)
Gallus gallus (Q5ZM71)
Xenopus tropicalis (A0A6I8QBE7)
Anolis carolinensis (G1KU86)
Danio rerio (E9QI95)
Oryzias latipes (H2LGG4)
Petromyzon marinus (XP_032812464.1)
Penaeus vannamei (XP_027222742.1)
Centruroides sculpturatus (XP_023243500.1)
Rhipicephalus sanguineus (XP_037507289.1)
Mizuhopecten yessoensis (XP_021366155.1)
Aplysia californica (XP_005097353.1)
Bombus terrestris (A0A6P5HNS2)
Drosophila melanogaster (Q2PDX8)
Thrips palmi (XP_034239953.1)
Komagataella phaffii (C4QZ05)
Saccharomyces cerevisiae (P50875)
similarity: 100% ≥80% ≥60% <60%
D
TAF12
vertebrate
invertebrate
yeast
Homo sapiens (Q16514)
Bos taurus (Q3T174)
Canis lupus (E2QVZ3)
Oryctolagus cuniculus (G1SKR2)
Mus musculus (Q8VE65)
Gallus gallus (Q5ZKX3)
Alligator mississippiensis (XP_019344041.1)
Tyto alba (XP_032858017.1)
Chelonia mydas (XP_037737490.1)
Anolis carolinensis (H9GH17)
Rhinatrema bivittatum (XP_029475403.1)
Pseudonaja textilis (XP_026558275.1)
Oryzias latipes (A0A3B3H6B2)
Danio rerio (Q6P013)
Xenopus tropicalis (F7EAB3)
Mizuhopecten yessoensis (OWF54813.1)
Bombus terrestris (A0A6P3U571)
Centruroides sculpturatus (XP_023233721.1)
Petromyzon marinus (XP_032834121.1)
Aplysia californica (XP_005101363.1)
Thrips palmi (XP_034253467.1)
Rhipicephalus sanguineus (XP_037519871.1)
Drosophila melanogaster (P49905)
Penaeus vannamei (XP_027217112.1)
Saccharomyces cerevisiae (Q03761)
Komagataella phaffii (C4R150)
linker (in yeast)
similarity: 100% ≥80% ≥60% <60%

**Fig. S7. Human versus yeast interactions between TRRAP/Tra1, SUPT20/Spt20 and TAF12.** (**A**) Location of the SUPT20H/Spt20 C-terminal region after superposition of human TRRAP and yeast Tra1. The C-terminal helix of yeast Spt20 aligns with helix one of the SUPT20H linker. (**B**) The sequence alignment of the SUPT20H/Spt20 C-terminal regions for 24 metazoan (SUPT20H) and two yeast (Spt20) species shows that the SUPT20H CTD is highly conserved in vertebrates, while it does not appear to exist in yeast. Secondary structure elements are indicated above the alignment. *: D291 forms a salt bridge with TRRAP R3746 (see Fig. S10H). Vertebrate and invertebrate sequences were pre-aligned to human SUPT20H, regions corresponding to the structured part in A were extracted and realigned with the yeast sequences corresponding to the region from helix 1 in A to the C-terminus. (**C**) Relative location of the TAF12 N-terminal region, based on the superposition shown in A. In yeast, an N-terminal linker of TAF12 wraps around the inside of the Tra1 FAT domain, while human TAF12 contacts TRRAP in a different location. The structured N-terminus of yeast TAF12 is located in the same relative position as the human SUPT20H CTD. (**D**) Zoomed-out sequence alignment of 24 metazoan and two yeast TAF12 subunits. Structured regions are colored as in C and the region corresponding to the linker in yeast is indicated. Aligned regions in B and D are colored by similarity in grey scale (annotated in D). In yeast, Taf12 contains a considerably longer N-terminus that appears to be unique to yeast. Sequences are labeled as: Scientific organism name (Uniprot or NCBI accession code). The organism selection corresponds to Table S1.

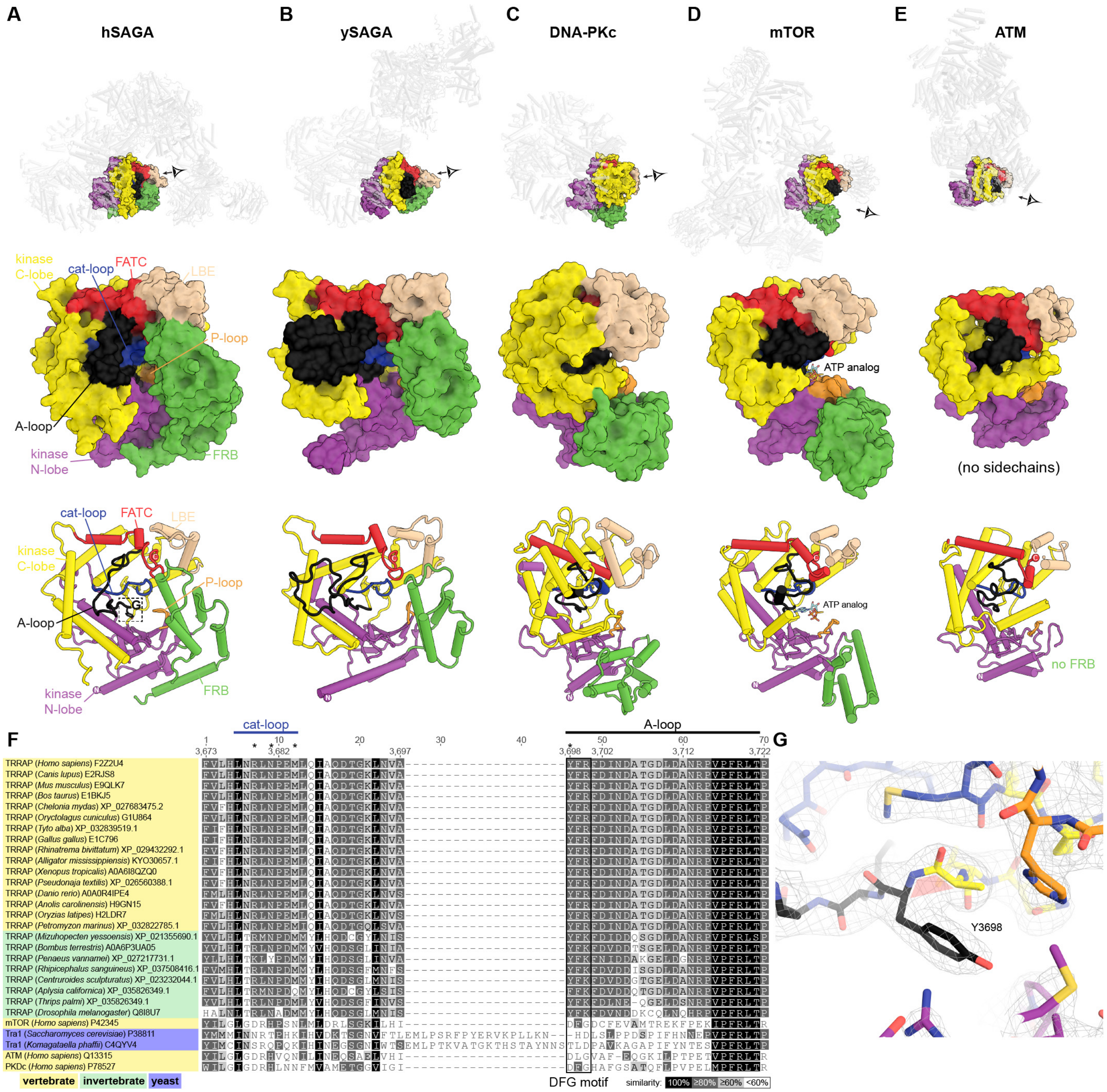

**Fig. S8. TRRAP ΨPI3K comparison and its integration into the HEAT repeat and FAT domain scaffold.** (**A-E**) ΨPI3K and its integration in hSAGA (A) and ySAGA(*15*) (B, PDB 6T9I), compared with the PI3K domains in DNA-PKc(*34*) (C, PDB 6ZFP), mTOR(*32*) (D, PDB 6BCX), and ATM(*35*) (E, PDB 5NP0). The panels below the overview show a close-up view as surface and cartoon representation of the active site entrance, as indicated in the top panels. Kinase elements are colored as indicated for hSAGA (FRB: FKBP-Rapamycin binding, P-loop: phosphate binding loop, LBE: mLST8 binding element, cat-loop: catalytic loop, FATC: FRAP-ATM-

TRRAP C-terminus, A-loop: activation loop). For mTOR the kinase crystal structure with ATPγS (*33*) (PDB 4JSP) is shown. In agreement with the comparison by Díaz-Santín *et al.* (*31*), the widely opened active site entrance in active kinases (C-E) is narrowed to a small tunnel in the pseudo-kinase in TRRAP (A) and Tra1 (B) by a rotation of the FRB domain. (**F**) Sequence alignment of the catalytic and activation loop region of twenty-nine ΨPI3Ks and PI3Ks, colored by similarity as indicated. *: residues proposed to be involved in catalysis in mTOR (*33*). (**G**) Residue Y3698 of hSAGA, equivalent to the first residue in the DFG motif of the activation loop in active kinases (*10*), adopts a cis-peptide bond, clearly defined by the density map (contoured at 9.0 σ).

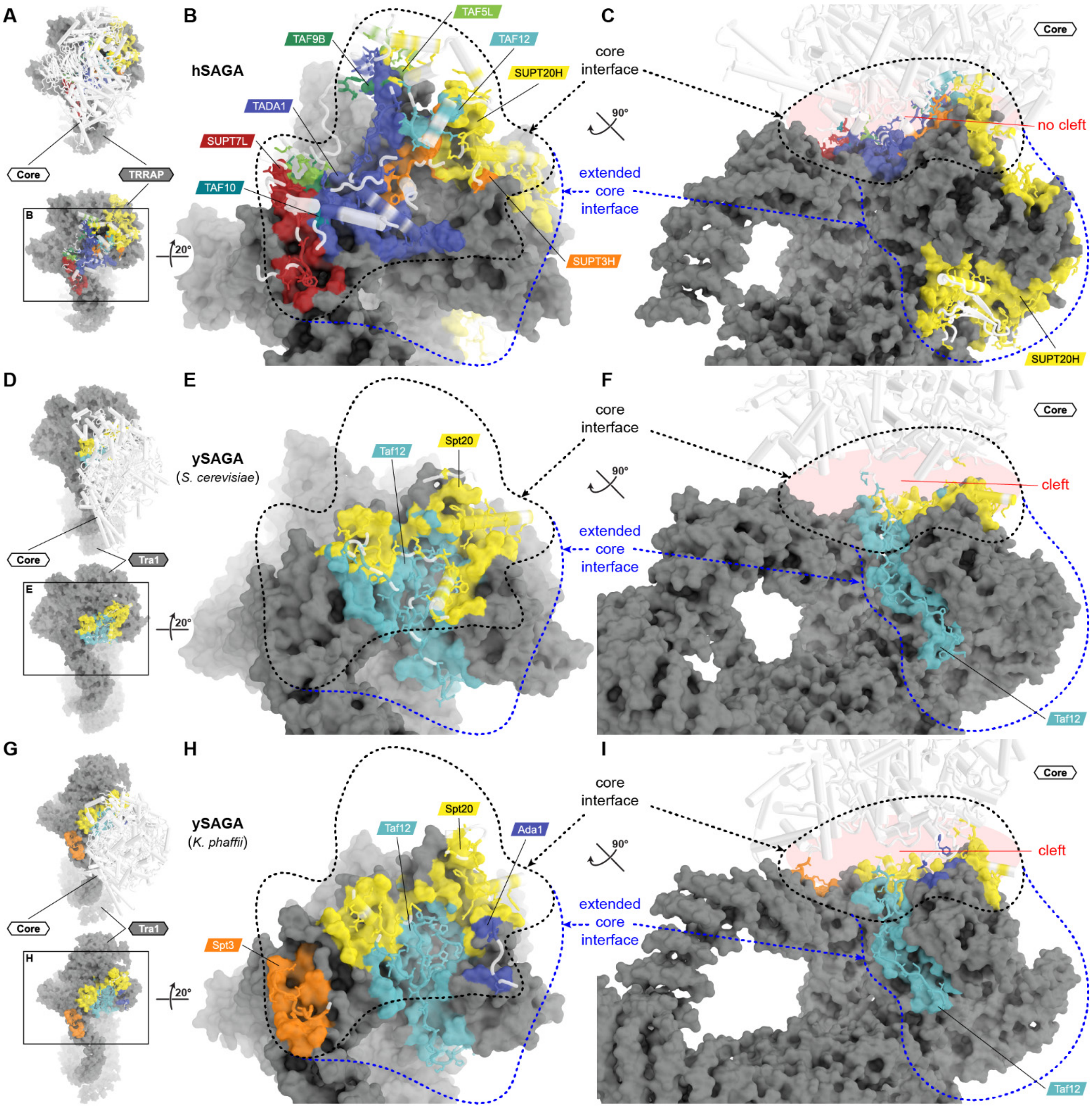

**J**

| TRRAP / Tra1 vs | all | **all**, except ~~SUPT20H~~ ~~Spt20~~ | **all**, except | **all**, except ~~SUPT20H~~ ~~Spt20~~ | SUPT20H / Spt20 | TAF12 / Taf12 | |
|---|---|---|---|---|---|---|---|
| **human**, PDB: 7KTR | 7,073 Å² | 2,661 Å² | 6,927 Å² | 2,514 Å² | 4,453 Å² | 179 Å² | |
| **yeast** (*S. cerevisiae*), PDB: 6T9I | 4,427 Å² | 2,880 Å² | 1,726 Å² | 0 Å² | 1,708 Å² | 2,862 Å² | complete |
| **yeast** (*K. phaffii*), PDB: 6TB4 | 4,217 Å² | 2,930 Å² | 1,960 Å² | 582 Å² | 1,370 Å² | 2,339 Å² | ≈ |
| **human**, PDB: 7KTR | 3,545 Å² | 2,661 Å² | 3,400 Å² | 2,514 Å² | 925 Å² | 179 Å² | |
| **yeast** (*S. cerevisiae*), PDB: 6T9I | 3,291 Å² | 1,744 Å² | 1,726 Å² | 0 Å² | 1,708 Å² | 1,726 Å² | in core interface |
| **yeast** (*K. phaffii*), PDB: 6TB4 | 3,532 Å² | 2,244 Å² | 1,960 Å² | 582 Å² | 1,370 Å² | 1,654 Å² | + |
| **human**, PDB: 7KTR | 3,528 Å² | 0 Å² | 3,528 Å² | 0 Å² | 3,528 Å² | 0 Å² | |
| **yeast** (*S. cerevisiae*), PDB: 6T9I | 1,152 Å² | 1,152 Å² | 0 Å² | 0 Å² | 0 Å² | 1,152 Å² | in extended core interface |
| **yeast** (*K. phaffii*), PDB: 6TB4 | 686 Å² | 686 Å² | 0 Å² | 0 Å² | 0 Å² | 686 Å² | |

**Fig. S9. Comparison of the core-TRRAP/Tra1 interface in human and yeast.** (**A**) Top view on the hSAGA core (white cartoon) and TRRAP module (grey surface). Parts of the core not in direct proximity to the interface have been removed in the lower panel. Interfacing residues on the TRRAP surface are colored based on their closest core subunit. Interfacing residues of core subunits are shown in stick representation and colored by subunit. (**B**) Magnified region from the box in A. The interface corresponding to the footprint of the core on the TRRAP surface is indicated with a black dashed outline (core interface). The interface created by extensions of core subunits that wrap around the TRRAP is indicated with a blue dashed outline (extended core interface). In hSAGA the latch and SUPT20H CTD (residues 274-428) contribute to the extended core interface. (**C**) Side view of the core and extended interface with the complete core shown as white cartoon. In contrast to ySAGA (F, I), hSAGA has no cleft between the core and TRRAP modules. (**D-F**) Same view as in A-C for *S. cerevisiae* ySAGA (PDB 6T9I) (*15*). (**G-H**) Same view as in A-C for *K. phaffii* ySAGA (PDB 6TB4) (*16*). (**F, I**) In both ySAGA structures the core and Tra1 modules are separated by a cleft. (**J**) Table with interface areas as indicated.

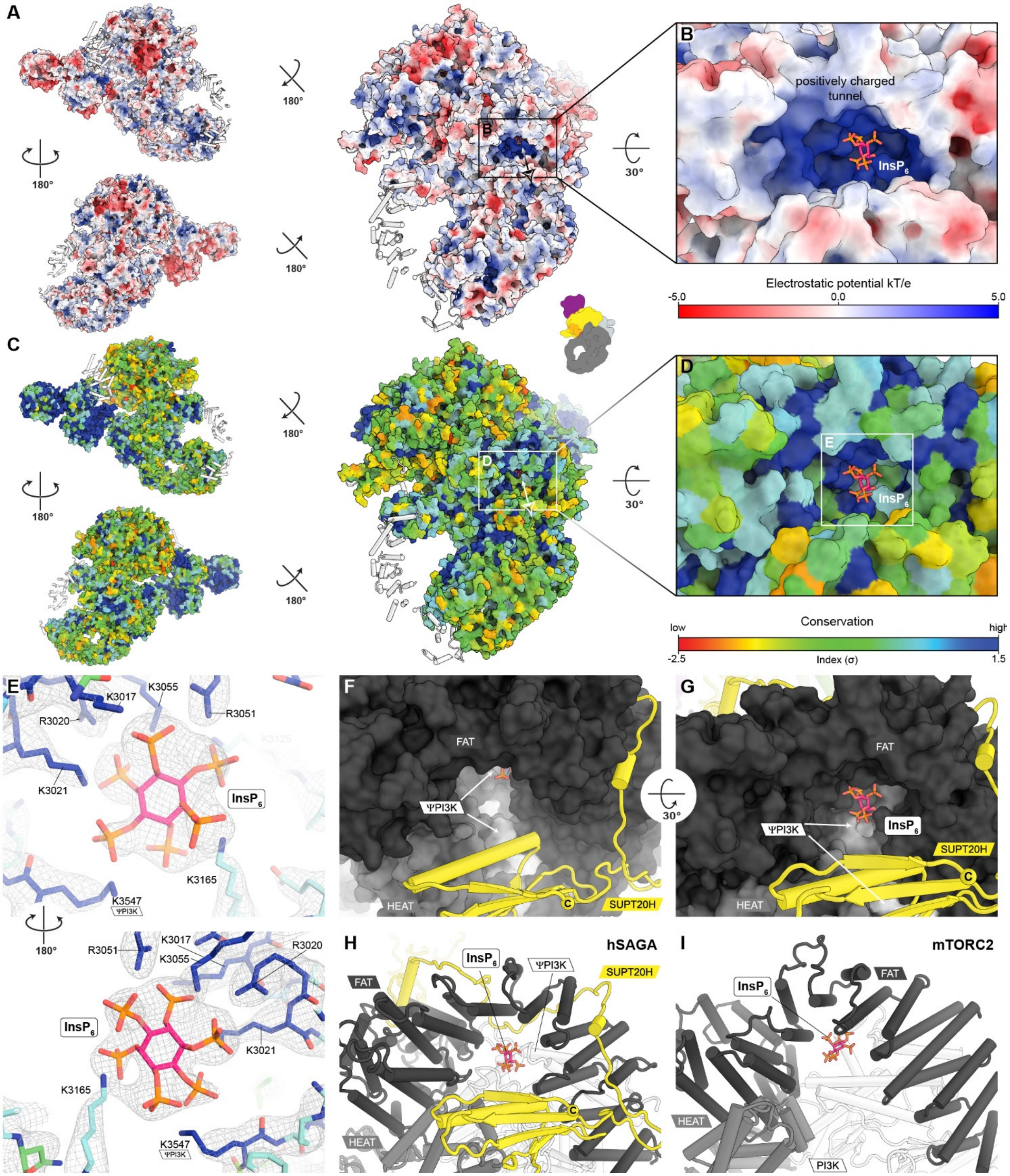

**Fig. S10. Electrostatic surface, conservation, and binding of inositol hexakisphosphate ($InsP_6$) to TRRAP.** (**A**) Electrostatic surface representation of hSAGA shown from three different views. Only regions with all-atom models have been included. Regions of lower sequence assignment confidence (UNKs) were excluded from the calculation and are shown as white cartoon. A highly positive charged tunnel between the FAT, HEAT, and ΨPI3K domains of TRRAP and SUPT20H is indicated. (**B**) Close-up view of the $InsP_6$ binding pocket within this tunnel. (**C**) Same views as in A colored by sequence conservation. Conservation was calculated based on sequence alignments of all the subunits in 24 metazoan SAGAs with the same subunit composition as hSAGA (see Table S1). (**D**) Close-up view of the $InsP_6$ binding pocket. $InsP_6$ (shown in stick representation) is bound by a ring of highly conserved residues. (**E**) Close-up view of the atomic model and the LocSpiral filtered multibody map (contoured at 11 $\sigma$) of TRRAP as indicated by the box in **D**. Residues involved in $InsP_6$ binding are indicated. All labeled residues are part of the TRRAP FAT domain except for K3547 (within ΨPI3K). Atoms are colored by conservation (carbon, see panel **D**), pink (carbon of $InsP_6$), red (oxygen), blue (nitrogen), or orange (phosphorus). (**F**) View as indicated by boxes in A and C; (**G**) view as in B and D (colored by domains and subunits). (**H, I**) Comparison of $InsP_6$ binding in hSAGA and in human mTORC2 (*46*) shown in cartoon representation. In both complexes $InsP_6$ binds in a similar location between the FAT and pseudo- and kinase domains of hSAGA and mTORC2, respectively.

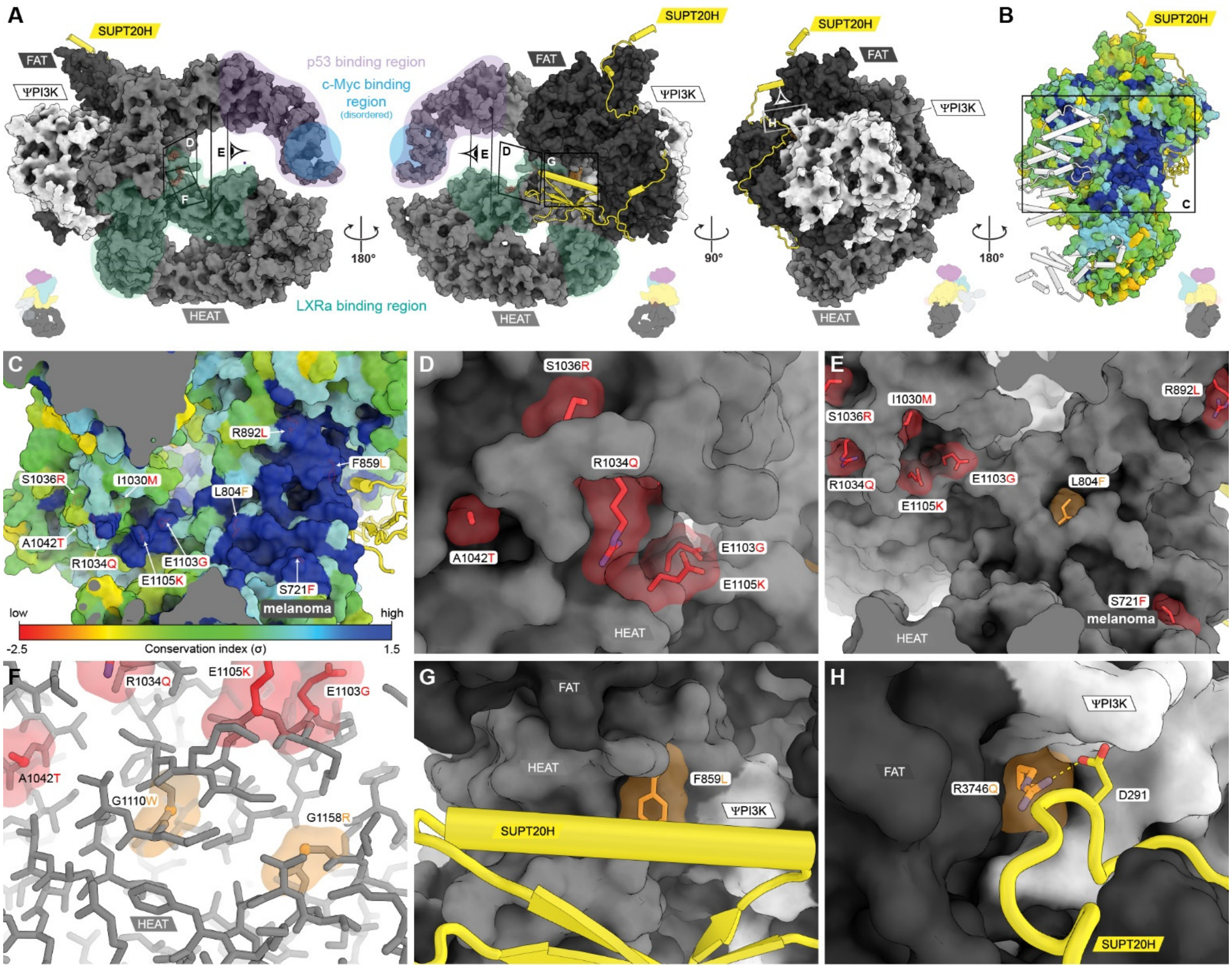

**Fig. S11. Activator binding regions and disease mutations in TRRAP.** (**A**) Surface representation of TRRAP colored by domains. Previously identified regions of activator binding are highlighted for p53 (*41*) (purple), c-Myc (*30*) (blue), and LXRa (*44*) (green). The sequence register assignment around the c-Myc binding region had low confidence and was modeled as UNKs. The c-Myc binding region is most likely located in a disordered loop between two helices of a HEAT repeat. Schematics at the bottom indicate the orientation relative to all modules. Boxes indicate the relative view in panels D-H. (**B**) Surface representation colored by conservation as depicted in Figure S10C. (**C**) Closeup of the region indicated with a box in B. Most disease mutations lie in a region of high sequence conservation. The location of the prevalent melanoma mutation S721F is indicated. (**D-H**) Close-up views of residues involved in cancer or identified as causative mutations in a study of patients with autism or syndromic intellectual disability (*42, 45*). Red residues are surface exposed and mutations might interfere with activator binding. Orange residues are buried, and mutations are likely to destabilize the structural integrity of TRRAP (F, not on the surface) or its interaction with SUPT20H (G, H). (**G**) F859L is located right at the interface with the SUPT20H CTD. (**H**) R3746 forms a salt bridge with D291 (see Fig. S7B) of the SUPT20H linker and its mutation (R3746Q) might destabilize the interaction with SUPT20H. The reported mutations from the literature for the canonical TRRAP isoform 1 (Uniprot Q9Y4A5) were mapped on the isoform observed in the structure (Uniprot F2Z2U4).

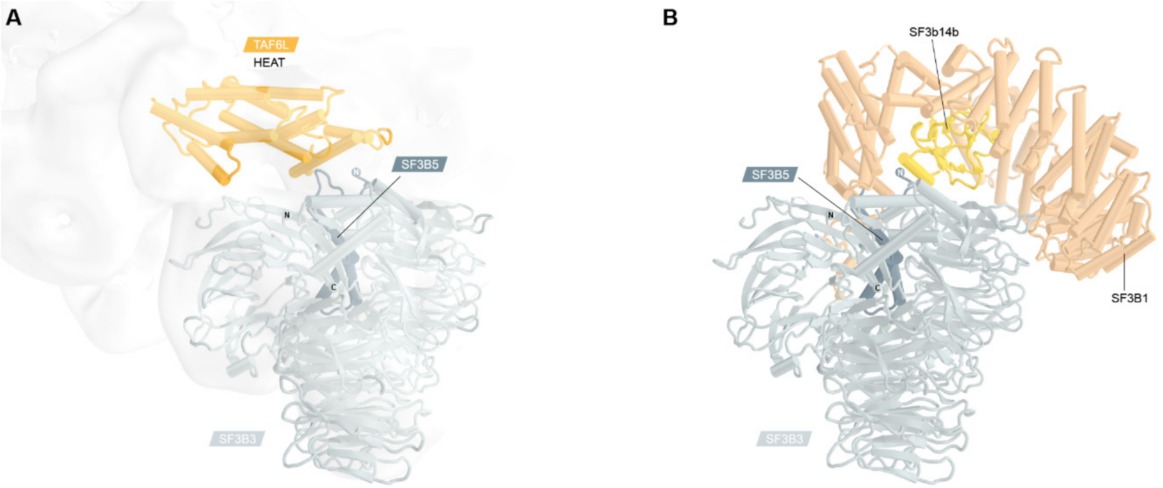

**Fig. S12. Comparison of the SF3B3/SF3B5 integration in hSAGA and the SF3b complex.** (**A**) The SPL module (SF3B3 and SF3B5 subunits) binds to the concave surface of the TAF6L HEAT domain. The negative stain map of hSAGA is shown in white transparency (contoured at 4.8 σ). (**B**) Crystal structure of the SF3b complex (*19*) (PDB 5IFE). The TAF6L HEAT domain of hSAGA is replaced by the SF3B1 HEAT repeat domain in the SF3b complex. Both domains share an overlapping binding region on the SF3B5 surface.

**Table S1. Sequence conservation of SAGA subunits in metazoan.**

| Common name | Scientific name | ATXN7 | ATXN7L3 | ENY2 | USP22 | KAT2A | SGF29 | TADA2B | TADA3 | SF3B3 | SF3B5 | SUPT20H | SUPT3H | SUPT7L | TADA1 | TAF10 | TAF12 | TAF5L | TAF6L | TAF9B | TRRAP | average |
|---|---|---|---|---|---|---|---|---|---|---|---|---|---|---|---|---|---|---|---|---|---|---|
| Human[m,v] | *Homo sapiens* | 100 | 100 | 100 | 100 | 100 | 100 | 100 | 100 | 100 | 100 | 100 | 100 | 100 | 100 | 100 | 100 | 100 | 100 | 100 | 100 | **100** |
| Mouse[m,v] | *Mus musculus* | 84 | 99 | 100 | 98 | 97 | 98 | 99 | 99 | 100 | 99 | 91 | 78 | 95 | 96 | 92 | 96 | 93 | 93 | 90 | 99 | **95** |
| Cattle[m,v] | *Bos taurus* | 89 | 99 | 100 | 97 | 98 | 99 | 83 | 100 | 100 | 100 | 86 | 80 | 97 | 97 | 93 | 99 | 94 | 94 | 85 | 99 | **94** |
| Dog[m,v] | *Canis lupus* | 93 | 97 | 95 | 98 | 98 | 95 | 23 | 100 | 100 | 100 | 82 | 68 | 98 | 99 | 97 | 99 | 99 | 94 | 94 | 99 | **91** |
| Rabbit[m,v] | *Oryctolagus cuniculus* | 91 | 97 | 78 | 97 | 94 | 91 | 23 | 99 | 97 | 100 | 82 | 91 | 96 | 98 | 95 | 99 | 99 | 89 | 95 | 97 | **90** |
| Alligator[v] | *Alligator mississippiensis* | 73 | 86 | 99 | 94 | 84 | 96 | 96 | 77 | 99 | 98 | 81 | 63 | 87 | 89 | 93 | 94 | 92 | 71 | 77 | 95 | **87** |
| Barn owl[v] | *Tyto alba* | 72 | 68 | 98 | 96 | 89 | 74 | 97 | 92 | 98 | 98 | 80 | 70 | 87 | 90 | 49 | 94 | 93 | 88 | 74 | 97 | **85** |
| Sea turtle[v] | *Chelonia mydas* | 36 | 86 | 98 | 94 | 87 | 97 | 96 | 92 | 99 | 99 | 80 | 67 | 88 | 88 | 64 | 93 | 93 | 71 | 79 | 97 | **85** |
| Snake[v] | *Pseudonaja textilis* | 66 | 86 | 98 | 91 | 82 | 97 | 93 | 89 | 99 | 97 | 73 | 67 | 85 | 89 | 81 | 90 | 90 | 64 | 75 | 93 | **85** |
| Lizard[v] | *Anolis carolinensis* | 72 | 81 | 98 | 89 | 82 | 96 | 94 | 92 | 99 | 97 | 67 | 66 | 86 | 84 | 51 | 92 | 92 | 68 | 76 | 90 | **84** |
| Two-lined caecilian[v] | *Rhinatrema bivittatum* | 63 | 84 | 99 | 94 | 69 | 96 | 90 | 92 | 97 | 97 | 72 | 66 | 79 | 70 | 63 | 91 | 86 | 68 | 78 | 96 | **83** |
| Chicken[v] | *Gallus gallus* | 69 | 86 | 96 | 92 | 84 | 95 | 21 | 93 | 99 | 98 | 71 | 67 | 87 | 84 | 67 | 95 | 92 | 59 | 79 | 97 | **82** |
| Japanese rice fish[v] | *Oryzias latipes* | 41 | 65 | 81 | 91 | 81 | 90 | 85 | 79 | 93 | 94 | 60 | 49 | 64 | 60 | 60 | 84 | 68 | 61 | 70 | 89 | **73** |
| Zebrafish[v] | *Danio rerio* | 42 | 68 | 85 | 92 | 83 | 91 | 25 | 78 | 93 | 94 | 62 | 57 | 70 | 65 | 61 | 79 | 67 | 63 | 74 | 92 | **72** |
| Frog[v] | *Xenopus tropicalis* | 4 | 11 | 94 | 92 | 7 | 96 | 8 | 88 | 97 | 95 | 70 | 66 | 79 | 81 | 64 | 76 | 81 | 58 | 77 | 94 | **67** |
| Lamprey[v] | *Petromyzon marinus* | 21 | 50 | 80 | 82 | 70 | 85 | 31 | 66 | 92 | 90 | 41 | 39 | 45 | 44 | 61 | 50 | 34 | 46 | 58 | 78 | **58** |
| Scallop[i] | *Mizuhopecten yessoensis* | 13 | 38 | 83 | 60 | 58 | 68 | 38 | 45 | 83 | 81 | 24 | 35 | 29 | 38 | 57 | 59 | 39 | 32 | 38 | 62 | **49** |
| Scorpion[i] | *Centruroides sculpturatus* | 14 | 39 | 73 | 57 | 61 | 65 | 41 | 46 | 84 | 76 | 21 | 20 | 30 | 40 | 58 | 49 | 31 | 30 | 46 | 56 | **47** |
| Tick[i] | *Rhipicephalus sanguineus* | 14 | 39 | 63 | 58 | 58 | 63 | 44 | 34 | 81 | 81 | 20 | 27 | 23 | 35 | 52 | 41 | 39 | 28 | 38 | 56 | **45** |
| Sea hare[i] | *Aplysia californica* | 13 | 32 | 79 | 57 | 54 | 56 | 37 | 40 | 81 | 78 | 23 | 32 | 24 | 33 | 58 | 38 | 35 | 23 | 41 | 56 | **45** |
| Bumblebee[i] | *Bombus terrestris* | 12 | 30 | 51 | 59 | 51 | 55 | 34 | 32 | 80 | 86 | 16 | 36 | 18 | 28 | 57 | 43 | 29 | 15 | 31 | 58 | **41** |
| Thrip[i] | *Thrips palmi* | 12 | 29 | 52 | 54 | 49 | 54 | 37 | 27 | 81 | 85 | 14 | 26 | 18 | 34 | 59 | 36 | 28 | 18 | 29 | 54 | **40** |
| Shrimp[i] | *Penaeus vannamei* | 12 | 30 | 59 | 18 | 54 | 48 | 43 | 37 | 79 | 76 | 23 | 32 | 24 | 28 | 55 | 33 | 29 | 16 | 34 | 57 | **39** |
| Fruit fly[i] | *Drosophila melanogaster* | 9 | 30 | 44 | 50 | 43 | 47 | 21 | 24 | 75 | 84 | 14 | 20 | 14 | 24 | 36 | 37 | 26 | 17 | 27 | 50 | **35** |
| Pichia pastoris[y] | *Komagataella phaffii* | 8 | 18 | 25 | 29 | 34 | 13 | 23 | 12 | N/A | N/A | 8 | 20 | 11 | 15 | 24 | 21 | 25 | 16 | 31 | 25 | **18** |
| Brewer's yeast[y] | *Saccharomyces cerevisiae* | 8 | 23 | 27 | 25 | 34 | 15 | 24 | 13 | N/A | N/A | 7 | 23 | 10 | 14 | 21 | 18 | 24 | 15 | 31 | 25 | **18** |

Sequence identities (%) and subunit names correspond to the human homologs and are sorted by decreasing average identity. N/A: not applicable (no known homolog). Vertebrates have an average sequence identity of ≥ 60%. Indices indicate classification into mammals (m), vertebrate (v), invertebrate (i), and yeast (y).

**Table S2. Mass spectrometric identification of hSAGA subunits present in the purified sample.**

| Accession Number | SAGA Subunit Name | SAGA module | Alternative Isoforms/Paralogs | Molecular Weight | Amino Acids | EUPC | Total Spectrum Count | PSM/AA* | Norm. Prop. to TRRAP |
|---|---|---|---|---|---|---|---|---|---|
| F2Z2U4 | TRRAP | TRRAP | Q9Y4A5, Q9Y4A5-2 | 438 kDa | 3859 | 299 | 771 | 0.1998 | 1.00 |
| Q92830 | KAT2A | HAT | Q92831 | 94 kDa | 837 | 57 | 176 | 0.2103 | 1.05 |
| Q15393 | SF3B3 | SPL | | 136 kDa | 1217 | 57 | 151 | 0.1241 | 0.62 |
| O75529 | TAF5L | Core | | 66 kDa | 589 | 53 | 171 | 0.2903 | 1.45 |
| Q8NEM7-3 | SUPT20H | Core | Q8NEM7-2, R4GND2 | 88 kDa | 811 | 40 | 134 | 0.1652 | 0.83 |
| O75528 | TADA3 | HAT | | 49 kDa | 432 | 32 | 108 | 0.25 | 1.25 |
| Q9Y6J9 | TAF6L | Core | | 68 kDa | 622 | 48 | 122 | 0.1961 | 0.98 |
| Q86TJ2 | TADA2b | HAT | | 48 kDa | 420 | 37 | 111 | 0.2643 | 1.32 |
| O15265 | ATXN7 | DUB | | 95 kDa | 892 | 35 | 106 | 0.1188 | 0.59 |
| Q96BN2 | TADA1 | Core | | 37 kDa | 335 | 29 | 90 | 0.2687 | 1.34 |
| Q9UPT9 | USP22 | DUB | | 60 kDa | 525 | 29 | 78 | 0.1486 | 0.74 |
| Q96ES7 | SGF29 | HAT | | 33 kDa | 293 | 19 | 62 | 0.2116 | 1.06 |
| Q9HBM6 | TAF9B | Core | Q16594 | 28 kDa | 251 | 20 | 63 | 0.251 | 1.26 |
| O94864 | SUPT7L | Core | O94864-2 | 46 kDa | 414 | 16 | 53 | 0.128 | 0.64 |
| Q12962 | TAF10 | Core | | 22 kDa | 218 | 10 | 40 | 0.1835 | 0.92 |
| Q14CW9 | ATXN7L3 | DUB | Q14CW9-2, Q5T6C5, Q9ULK2 | 39 kDa | 347 | 17 | 47 | 0.1354 | 0.68 |
| O75486 | SUPT3H | Core | O75486-4, Q5U608 | 36 kDa | 317 | 11 | 36 | 0.1136 | 0.57 |
| Q16514 | TAF12 | Core | Q16514-2 | 18 kDa | 161 | 9 | 21 | 0.1304 | 0.65 |
| Q9BWJ5 | SF3B5 | SPL | | 10 kDa | 86 | 7 | 25 | 0.2907 | 1.45 |
| Q9NPA8 | ENY2 | DUB | Q9NPA8-2 | 12 kDa | 101 | 5 | 17 | 0.1683 | 0.84 |

Proteins are shown as identified only if the protein probability is >99% and the peptide threshold >95%. Paralogs present or isoforms indistinguishable by MS are listed in Column 4. If multiple isoforms were indistinguishable by MS, the modeled isoform is listed in Column 1. EUPC = Exclusive Unique Peptide Count. *PSM/AA equals Total Spectrum Count divided by # Amino Acids. Column 10 shows PSM/AA values normalized to TRRAP=1 to show the approximate stoichiometry of each subunit. The table shows only subunits of the hSAGA complex. For the full list of peptides recovered, see Table S3.

**Table S3. Mass spectrometric identification of all proteins in the purified sample.**

| Total Spectrum Count | SAGA Subunit Name | Accession Number | Alternative Isoforms |
|---|---|---|---|
| 771 | TRRAP | Q9Y4A5 | Q9Y4A5-2 |
| 479 | TRRAP | F2Z2U4 | |
| 176 | KAT2A | Q92830 | |
| 171 | TAF5L | O75529 | |
| 151 | SF3B3 | Q15393 | |
| 134 | SUPT20H | Q8NEM7-3 | Q8NEM7-2, R4GND2 |
| 122 | TAF6L | Q9Y6J9 | |
| 111 | TADA2b | Q86TJ2 | |
| 108 | TADA3 | O75528 | |
| 106 | ATXN7 | O15265 | |
| 90 | TADA1 | Q96BN2 | |
| 78 | USP22 | Q9UPT9 | |
| 65 | KAT2B | Q92831 | |
| 63 | TAF9B | Q9HBM6 | |
| 62 | SGF29 | Q96ES7 | |
| 57 | ATXN7L1 | Q9ULK2 | A4D0Q3 |
| 53 | SUPT7L | O94864 | O94864-2 |
| 51 | ATXN7L2 | Q5T6C5 | |
| 49 | | A0A3B3ITZ9 | Q9Y2W1 |
| 47 | ATXN7L3 | Q14CW9 | Q14CW9-2 |
| 43 | TAF9 | Q16594 | |
| 40 | TAF10 | Q12962 | |
| 36 | SUPT3H | O75486 | O75486-4, Q5U608 |
| 29 | | A0A4D5RAC7 | A0A4D5RAC9, P68104, Q53G85, Q53GE9, Q53HQ7, Q53HR5, Q6IPN6, QPIPT9 |
| 27 | | A0A1W2PPS1 | B4DLR3, Q00839, Q00839-2 |
| 25 | SF3B5 | Q9BWJ5 | |
| 22 | | Q1KMD3 | |
| 21 | TAF12 | Q16514 | Q16514-2 |
| 19 | | P60709 | P63261, Q53G76, Q53GK6 |
| 17 | ENY2 | Q9NPA8 | Q9NPA8-2 |
| 17 | | P84090 | |
| 13 | | P11142 | P11142-2, Q53HF2 |
| 12 | | E9PK91 | Q9NYF8, Q9NYF8-2, Q9NYF8-3 |
| 10 | | Q8NF21 | |
| 10 | | B4DE59 | |
| 5 | | Q14966 | Q14966-3 |
| 5 | | Q9Y3Y2 | Q9Y3Y2-3, Q9Y3Y2-4, X6R700 |
| 5 | | A0A0G2JIW1 | A0A1U9X7W4, B4DFN9, P0DMV8, P0DMV8-2, Q59EJ3 |
| 5 | | B2RBD5 | |
| 4 | | Q6UWP8 | |
| 4 | | I0B0K3 | I0B0K4, I0B0K5, I0B0K6, I0B0K7, I0B0K8, P20930, Q05331 |
| 4 | | P67809 | |
| 4 | | P31151 | |
| 3 | | M0R0R2 | P46782 |
| 3 | | P06702 | |
| 3 | | P04406 | P04406-2, Q0QET7, Q2TSD0 |
| 3 | | Q53GA7 | |
| 3 | | A6NMY6 | |
| 2 | | P11021 | |
| 2 | | C9J352 | C9J4U1, C9JWD9, C9JYI4, F2Z3G4, P52294, Q5BKZ2 |
| 2 | | Q05DU1 | Q05DU1, Q9NVN8 |

Proteins are shown as identified only if the protein probability is >99%, the peptide threshold >95%, and at least 2 peptides were found. Isoforms that could not be distinguished by spectra are listed under "Alternative isoforms".

**Table S4. Cryo-electron microscopic data collection and refinement statistics**

| | Cryo-EM | Negative stain |
|---|---|---|
| **Data collection and processing** | | |
| Microscope | FEI Titan Krios G2 | FEI Technai F20 |
| Camera | Gatan K3 Summit (super resolution) | Gatan UltraScan4000 |
| Voltage (keV) | 300 | 120 |
| Magnification | 64,000 | 80,000 |
| Defocus range(μm) | 0.9-3.4 | 0.4-3.9 |
| Micrographs / Movies | 10,224 | 1 |
| Frames per movie | 50 | N/A |
| Pixel size (Å) | 1.187 | 1.4 |
| Total dose (e- $Å^{-2}$) | 50 | 35 |
| Particles initial / final | 3,167,367 / 357,441 | 47,790 / 3,157 |
| Map resolution (Å) | 2.9 | 19 |
| **Refinements** | | |
| Method | real space, adp | rigid body* |
| C-beta deviations | 0.00 | 0.00 |
| Rotamer outliers (%) | 0.02 | 0.02 |
| All-atom Clashscore | 3.42 | 4.79 |
| MolProbity score | 1.50 | 1.62 |
| Ramachandran Plot (outliers / favored) (%) | 0.04 / 94.72 | 0.05 / 94.57 |
| Rama-Z score, whole (r.m.s. Rama-Z) | 0.16 (0.12) | -0.35 (0.11) |
| Map-to-model cross-correlation | 0.80 | 0.52 |
| Non-hydrogen atoms | 40,337 | 51,173 |
| Ligand / Non-hydrogen atoms | 36 | 0 |
| Protein residues | 5,169 | 6,632 |
| Mean B-factor (protein / ligand) ($Å^2$) | 73.6 / 66.6 | not refined |
| R.m.s. deviations | | |
| Bond lengths (Å) | 0.004 | 0.005 |
| Bond angles (º) | 0.62 | 0.99 |

*: Three bodies were fit comprising the SF3B3/SF3B5 subunits of SF3b (*19*) (PDB 5IFE), a homology model based on human TAF6 (*5*) (PDB 6MZL), and the cryo-EM structure of hSAGA. adp: atomic displacement parameters.

**Movie S1. Overview of the negative stain and cryo-EM reconstructions as well as the atomic model of hSAGA.**
